# An ancient role for the *CYP73* gene family in *t*-cinnamic acid 4-hydroxylation, phenylpropanoid biosynthesis and embryophyte development

**DOI:** 10.1101/2023.08.20.551634

**Authors:** Samuel Knosp, Lucie Kriegshauser, Kanade Tatsumi, Ludivine Malherbe, Gertrud Wiedemann, Bénédicte Bakan, Takayuki Kohchi, Ralf Reski, Hugues Renault

## Abstract

The phenylpropanoid pathway is a plant metabolism intimately linked to the transition to terrestrial life. It produces phenolic compounds that play essential roles in stress mitigation and ecological interactions. The pathway also provides the building blocks for hydrophobic polymers that form apoplastic diffusion barriers and make up a significant fraction of the land plant biomass. Despite its significance in embryophytes (i.e., land plants), the origin and evolutionary history of the phenylpropanoid pathway remain poorly understood. In particular, little is known about the organization and function of the pathway in bryophytes, the non-vascular embryophytes. In this study, we conducted a multidisciplinary analysis of the *CYP73* gene family that encodes *t*-cinnamic acid 4-hydroxylase (C4H), the first plant-specific enzyme in the pathway. Our results indicate that C4H activity originated with the emergence of the *CYP73* gene family in an ancestor of land plants and was supported by an arginine residue that stabilizes its substrate in the active site. C4H deficiency in the moss *Physcomitrium patens*, the liverwort *Marchantia polymorpha* and the hornwort *Anthoceros agrestis* resulted in a shortage of phenylpropanoids and abnormal plant development. The latter could be rescued in the moss by the exogenous supply of *p*-coumaric acid, the product of C4H. Our findings establish the emergence of the *CYP73* gene family as a foundational event for the development of the canonical plant phenylpropanoid pathway and underscores the deep-rooted conservation of the C4H enzyme function in embryophyte biology.

## INTRODUCTION

Green plants (i.e., *Viridiplantae*) have evolved over approximately one billion years and have thrived wherever light was available thanks to their photosynthetic capabilities. The successful evolutionary history of *Viridiplantae* have yielded an astonishing diversity of forms – from unicellular marine algae to giant redwood trees – and propelled them as the most prevalent living group on Earth from a biomass standpoint (1).

Most of this biomass is found on land (1), reflecting the cornerstone functions performed by plants in terrestrial ecosystems, primarily as an entry point for solar energy. Recent phylogenetic evidences indicate that modern land plants, also known as embryophytes, have a unique origin and emerged from freshwater algae about half a billion years ago (2, 3). The radical change in habitat experienced by early embryophytes, an evolutionary milestone called terrestrialization, involved profound morpho-physiological adaptations. In particular, the ability to synthesize a wide range of metabolites was instrumental by mitigating effects of terrestrial constraints (e.g., UV, lack of buoyancy, drought) and by providing chemical mediators for fast-expanding ecological interactions.

One of the most iconic plant metabolisms associated with terrestrialization is the phenylpropanoid pathway, which generates a suite of phenolic compounds that effectively address terrestrial challenges (4, 5). This pathway leads to the synthesis of widespread polyphenolic molecules (e.g., flavonoids) and phenolic esters/amides (e.g., chlorogenic acids), which act as powerful UV screens and antioxidants. The phenylpropanoid pathway also supplies precursors of the four hydrophobic polymers cutin, suberin, sporopollenin and lignin that strengthen and waterproof the cell wall. These polymers form the framework of various apoplastic diffusion barriers (e.g., cuticle, pollen coat, Casparian strip) and thus help plants to shield their tissues from external aggressions, and to manage water and solutes efficiently. In accordance to these essential functions, hydrophobic biopolymers make up a significant fraction of embryophyte biomass. Lignin for instance is regarded as the second most abundant biopolymer on the planet after cellulose, accounting for *ca*. 30% of biosphere organic carbon (6).

In order to support high biomass production and multiple physiological functions, the phenylpropanoid pathway is meticulously organized and regulated (7, 8). The first three steps of this pathway, collectively known as the general phenylpropanoid pathway (**Fig. 1A**), are obligatory and hence process the entire metabolic flux. The initial step involves the deamination of phenylalanine by the phenylalanine ammonia-lyase (PAL) to produce *trans*-cinnamic acid, which is the first *bona fide* phenylpropanoid molecule. The phenolic ring of *t*-cinnamic acid is then hydroxylated at position 4 (*para* position) by the cinnamic acid 4-hydroxylase (C4H) to generate *para*-coumaric acid. Next, *p*-coumaric acid is activated with coenzyme A by 4-coumarate:CoA ligase (4CL), which leads to the formation of *p*-coumaroyl-CoA, a branch-point molecule that feeds into the various downstream pathways (**Fig. 1A**).

**Figure 1.**
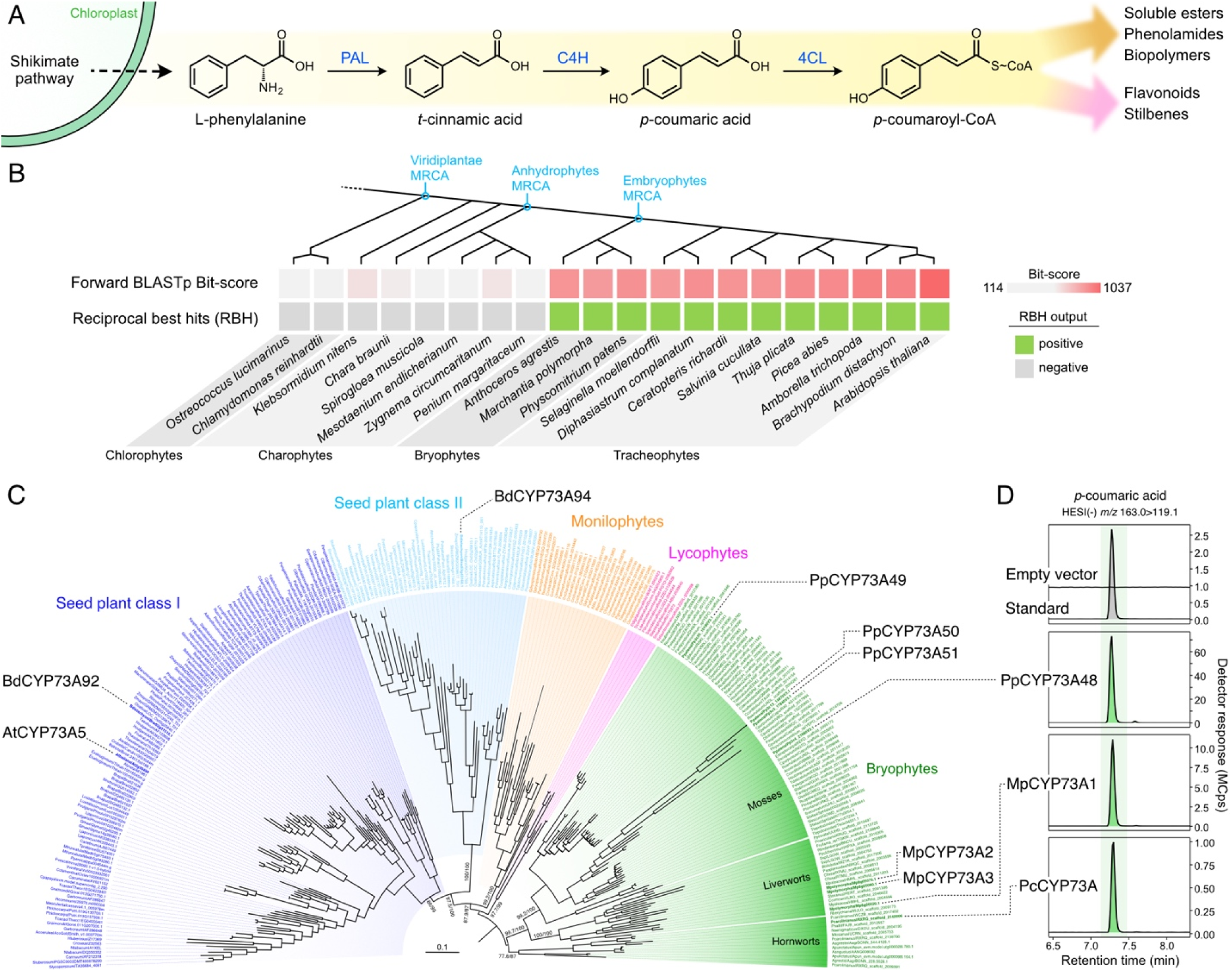
Evolutionary history of the *CYP73* family encoding cinnamic acid 4-hydroxylase. (A) Schematic representation of the three steps of the general phenylpropanoid pathway. PAL, phenylalanine ammonia-lyase; C4H, cinnamic acid 4-hydroxylase; 4CL, 4-coumarate:CoA ligase. (B) Search for *AtCYP73A5* orthologs by reciprocal best hit (RBH) strategy. MRCA, most recent common ancestor. (C) ML nucleotide tree (IQ-TREE2, GTR+F+I+R7) describing the phylogenetic relationships between 275 *CYP73* homologous sequences. SH-aLRT test and ultrafast bootstrap (1,000 pseudo-replicates each) supports are indicated on main branches. CYP73 homologs relevant to the present study are indicated. At, *Arabidopsis thaliana*; Bd, *Brachypodium distachyon;* Mp, *Marchantia polymorpha*; Pc, *Phaeoceros carolinianus*. Scale bar represents the number of nucleotide substitutions per site. Tree was rooted according to the midpoint method. (D) Representative UHPLC-MS/MS chromatograms showing the production *in vitro* of *p*-coumaric acid from *t*-cinnamic acid (C4H activity) by candidate CYP73 proteins from the three major bryophyte groups: mosses (PpCYP73A48), liverworts (MpCYP73A1) and hornworts (PcCYP73A). Assays performed with microsomes derived from yeast transformed with an empty vector were used as negative control.

PAL activity is not exclusive to plants and is also found in cyanobacteria and fungi (9), thus making C4H the first plant-specific step in the phenylpropanoid pathway (10). C4H enzymes belong to the family 73 of cytochrome P450 monooxygenases (CYP73). Cytochromes P450 are highly diversified enzymes that catalyze irreversible and often rate-determining reactions and, as such, are considered to be key drivers of plant metabolism evolution and diversification, and control points in metabolic pathways (11–13). Accordingly, inactivation of the single *CYP73* gene from the tracheophyte model *Arabidopsis thaliana* led to dramatic developmental defects and phenylpropanoid deficiency (14), pointing to the critical role of C4H in tracheophyte physiology. No such *in planta* investigation has been performed to date in a bryophyte model although these data appear key to improve our understanding of the evolution and early functions of C4H and, by extension, of the phenylpropanoid pathway as a whole.

Here we report a multidisciplinary study of C4H-encoding *CYP73* genes in representative species of the three major bryophyte groups. We show that C4H activity emerged with the rise of the *CYP73* family in an embryophyte ancestor and identify key residues supporting its catalytic activity. We further show that *CYP73* deficiency in the moss *Physcomitrium patens* annihilates the ability to produce phenylpropanoid and strongly alters plant development. Such a function was confirmed in the liverwort *Marchantia polymorpha* and the hornwort *Anthoceros agrestis*, indicating a pivotal and conserved role of *CYP73* in embryophyte physiology.

## RESULTS

### CYP73-catalyzed C4H activity emerged in an embryophyte progenitor

Thus far, the only known enzymes capable of catalyzing the 4-hydroxylation of *t*-cinnamic acid (C4H activity) are the cytochromes P450 monooxygenases from the 73 family (CYP73). To investigate the origin of C4H, a survey of *CYP73* genes was conducted using a reciprocal best hits approach in 20 *Viridiplantae* genomes, including recent charophyte genomes (**Fig. 1B**, **Tab. S1**). The results revealed that potential *CYP73* orthologs were only found in embryophytes, suggesting that the origin of this CYP family can be traced back to a progenitor of embryophytes. The charophyte *Klebsormidium nitens* had the best hit among non-embryophyte plants, but it shared greater homology with the CYP98 family rather than with CYP73 (**Tab. S1**). Accordingly, an *in vitro* enzyme assay with the corresponding *K. nitens* recombinant protein did not result in detectable C4H activity under the tested conditions (**Fig. S1**).

The evolution of the *CYP73* family was further studied by reconstructing the phylogenetic relationships between 275 *CYP73* sequences, which encompassed the entire range of embryophyte diversity, including 85 bryophyte homologous sequences. Aside the previously reported duplication that occurred in a seed plant ancestor (10), the topology of the tree reflected embryophyte systematics (**Fig. 1C**). *CYP73* family was found fairly diversified in bryophytes with for instance four paralogs in the moss *Physcomitrium patens* (*PpCYP73A48-51*) and three in the liverwort *Marchantia polymorpha* (*MpCYP73A1-3*). In mosses, this diversity could be at least in part explained by an early duplication that gave rise to two major *CYP73* groups (**Fig. 1C**). This duplication occurred less than 350 million year ago, after the *Takakia* and *Sphagnum* genera branched off (15). Next, we performed C4H assays with yeast microsomes containing recombinant CYP73 proteins from *P. patens* (PpCYP73A48), *M. polymorpha* (MpCYP73A1) and the hornwort *Phaeoceros carolinianus* (PcCYP73A). The three bryophyte CYP73 proteins catalyzed the production of *p*-coumaric acid *in vitro*, whereas microsomes derived from yeast transformed with the empty vector did not (**Fig. 1D**). In combination with biochemical data previously obtained in tracheophytes (7, 10), these results indicate that C4H activity is conserved in embryophytes and emerged in a progenitor of embryophytes with the rise of the *CYP73* family.

### C4H activity is supported by a conserved arginine residue

To gain a deeper understanding of the origin of C4H activity, we investigated the underlying structural determinants. We employed homology modeling to construct the three-dimensional structure of the *P. patens* CYP73A48 enzyme, which had the ability to catalyze C4H activity *in vitro* (**Fig. 1D**), based on the crystal structure of *Sorghum bicolor* C4H (16). After docking the heme prosthetic group into the PpCYP73A48 structure, we went on with docking *t*-cinnamic acid into its active site. The lowest Gibbs free energy pose (-6.7 kcal/mol) predicted that position 4 of the *t*-cinnamic acid phenolic ring was facing heme at a distance of 4.3 Å (**Fig. 2A**), which is consistent with C4H activity. The docking experiment also identified three residues in the F helix of PpCYP73A48 – arginine 225, serine 226, and glutamine 230 – presumably able to form hydrogen bonds with the carboxylic function of *t*-cinnamic acid and to hold it in the proper orientation with respect to the heme reaction center (**Fig. 2A**). Examination of CYP73 multiple protein alignment and corresponding weblogo showed that these three residues were highly conserved across the 275 sequences used for phylogenetic analysis (**Fig. 2B**).

**Figure 2.**
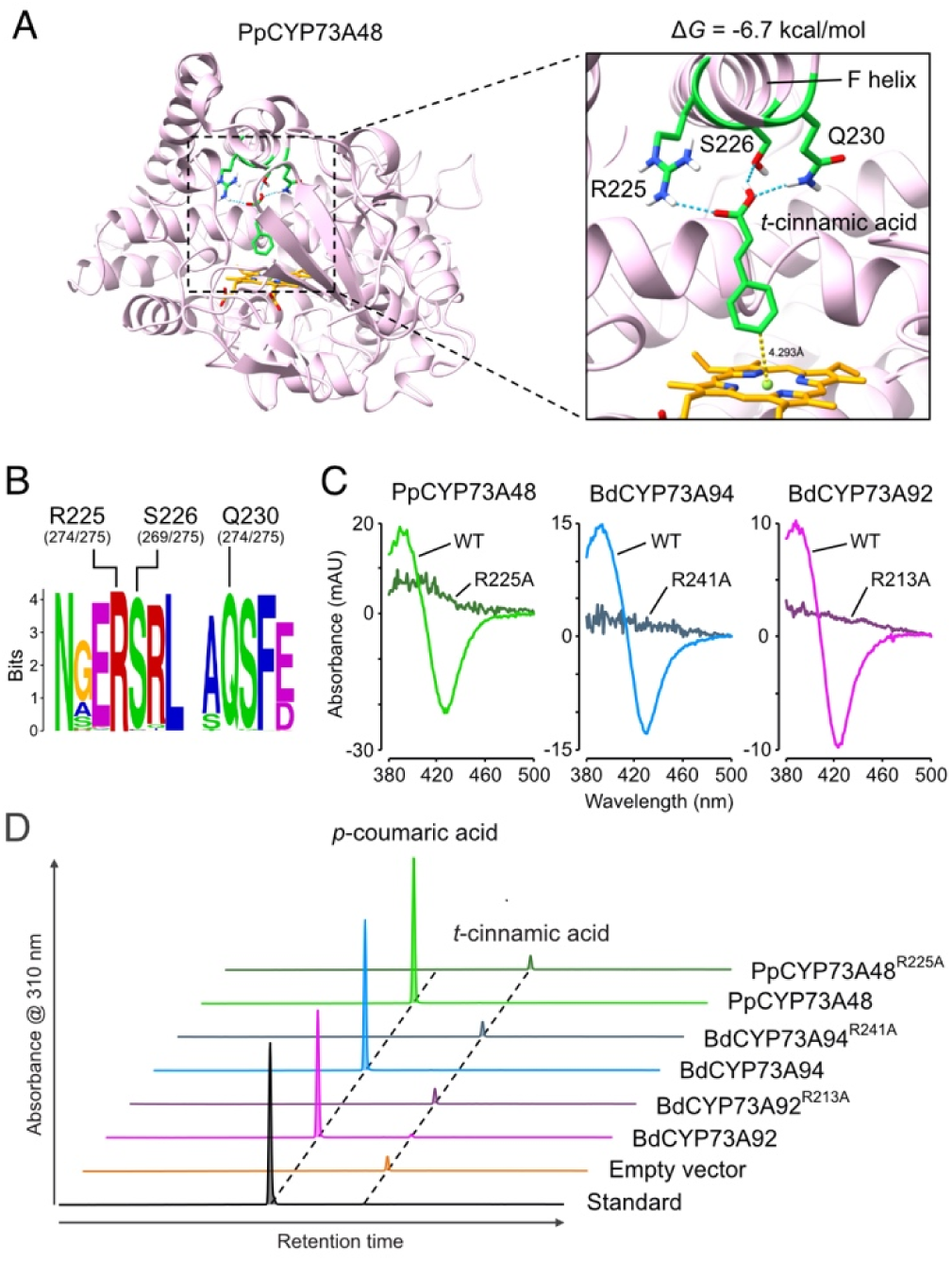
Structural determinants of cinnamic acid 4-hydroxylase activity in CYP73. (A) Docking of *t*-cinnamic acid in the active site of the *P. patens* CYP73A48 enzyme. Pose corresponding to the lowest Gibbs free energy (ΔG) predicts that R225, S226 and Q230 residues can establish hydrogen bonds with the carboxyl group of *t*-cinnamic acid. In this configuration, *para* position (or position 4) of the phenolic ring faces the heme prosthetic group at a distance of 4.3 Å. (B) Weblogo showing the conservation of R225, S226 and Q230 residues across the 275 CYP73 sequences used for the phylogenetic analysis. For each residue, absolute count is provided between brackets. (C) Representative type I difference spectra showing the loss of cinnamic acid binding ability of *P. patens* CYP73A48, *B. distachyon* CYP73A94 and *B. distachyon* CYP73A92 enzymes mutated in the R225 residue, or its equivalent. Positively charged arginine residue was replaced with neutral alanine. (D) Representative HPLC-UV chromatograms of *in vitro* enzyme assays indicating loss of C4H activity in CYP73 enzymes mutated in the R225 residue, or its equivalent.

To confirm the predictions of docking experiments, we focused on the R225 residue of PpCYP73A48, which has a positive charge and can thus strongly interact with the negatively charged carboxyl group of *t*-cinnamic acid. We expanded the evolutionary scope of the experiment by including the previously characterized and phylogenetically distant CYP73A94 and CYP73A92 proteins from *Brachypodium distachyon* (10) (**Fig. 1C**). We replaced the R225 residue, or its equivalent in BdCYP73A94 (R241) and BdCYP73A92 (R213), with a non-polar and uncharged alanine residue. Carbon monoxide-induced difference spectra showed that the R>A substitution did not affect overall the structural stability of CYP73 proteins (**Fig. S2**). On the contrary, substrate-induced type I difference spectrum revealed that all three R>A mutated CYP73 proteins lost their ability to bind to *t*-cinnamic acid compared to their wild-type counterparts (**Fig. 2C**). Consequently, the R>A CYP73 mutant proteins did not produce *p*-coumaric acid from *t*-cinnamic acid in *in vitro* assays (**Fig. 2D**), indicating that the evolutionarily conserved R225 residue was critical for C4H activity.

### *CYP73* deficiency strongly alters moss development

We performed in planta investigation of the *CYP73* genes identified in the moss *P. patens* (**Fig. 1C**). Scrutiny of public RNA-seq data revealed that only three paralogs were expressed in at least one condition across *P. patens* tissues, with *PpCYP73A48* and *PpCYP73A49* being the most prominent (**Fig. 3A**). We therefore decided to disregard the *PpCYP73A50* paralog, as it displayed no expression and also exhibited substantial variation among the protein region containing the three residues interacting with *t*-cinnamic acid (**Fig. 3B**). We then performed *in vitro* assays with PpCYP73A48, PpCYP73A49 and PpCYP73A51 recombinant proteins tagged with 3xHA. While all three proteins were confirmed to be present in yeast microsomes through Western blot analysis (**Fig. S3**), only PpCYP73A48-3xHA and PpCYP73A49-3xHA demonstrated the ability to convert *t*-cinnamic acid into *p*-coumaric acid (**Fig. 3C**). This observation was further supported in planta by the fact that *PpCYP73A48* and *PpCYP73A49* fully complemented the stunted phenotype of the Arabidopsis *cyp73a5-*1 mutant (**Fig. 3D**), whereas no mutant plant rescued with *PpCYP73A51* was ever found.

**Figure 3.**
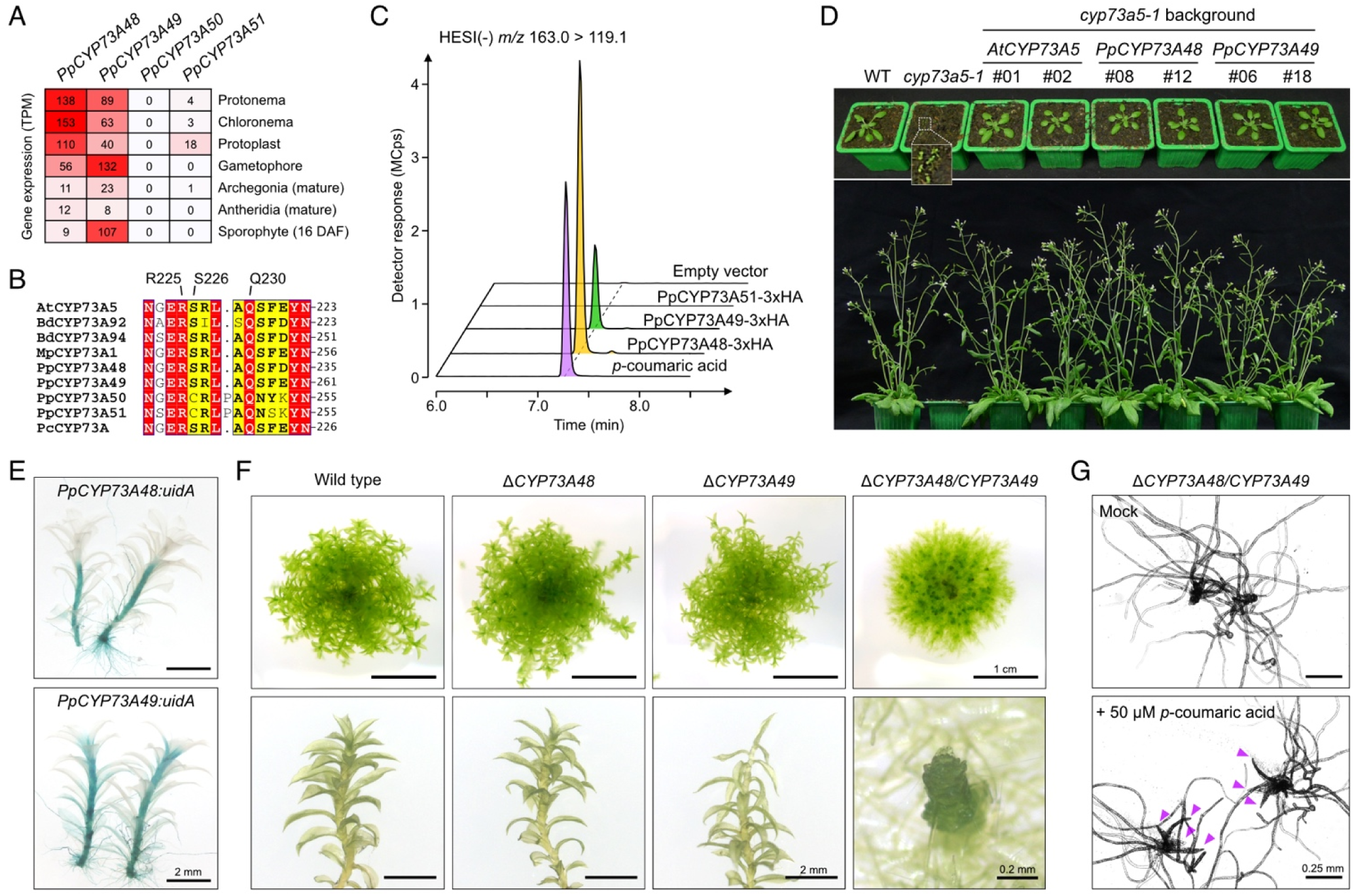
Functional analysis of *CYP73* genes from the moss *Physcomitrium patens*. (A) Expression profiles of the four *P. patens CYP73* paralogs in various tissues. Data are derived from the CoNekT database (65). DAF, days after fertilization; TPM, Transcripts Per Kilobase Million. (B) Angiosperm and bryophyte CYP73 multiple-sequence alignment centered on the protein region encompassing cinnamic acid binding residues identified by docking. At, *Arabidopsis thaliana*, Bd, *Brachypodium distachyon*; Mp, *Marchantia polymorpha*; Pc, *Phaeoceros carolinianus*; Pp, *Physcomitrium patens*. (C) Representative UHPLC-MS/MS chromatograms of *in vitro* C4H assays performed with recombinant PpCYP73-3xHA tagged proteins. Assays performed with microsomes derived from yeast transformed with an empty vector were used as negative control. (D) Phenotype of *Arabidopsis thaliana* three-week-old (upper panel) and six-week-old (lower panel) wild type, *cyp73a5-1* mutant and *cyp73a5-1* complemented with *AtCYP73A5*, *PpCYP73A48* and *PpCYP73A49* coding sequences. Two independent complemented lines are depicted for each gene. (E) Representative GUS staining pattern in two-month-old gametophores of *PpCYP73A48:uidA* and *PpCYP73A49:uidA* reporter lines. (F) Pictures of two-month-old colonies of wild type, Δ*PpCYP73A48* and Δ*PpCYP73A49* single mutants, and Δ*PpCYP73A48/CYP73A49* double mutant. Close-ups on individual gametophores from each genotype are visible in the lower part. (G) Pictures of five-week-old Δ*PpCYP73A48/CYP73A49* gametophores grown in low-melting point agarose Knop medium supplemented, or not (control), with 50 µM *p*-coumaric acid. Magenta arrowheads point to developed phyllids.

To refine the analysis of *PpCYP73A48* and *PpCYP73A49* expression patterns, we generated knock-in lines where the STOP codon was replaced with the *uidA* reporter gene. These lines revealed that *PpCYP73A48* and *PpCYP73A49* had overlapping expression domains in gametophores, particularly in the stem (**Fig. 3E**). It was noted however that *PpCYP73A49* expression extended further to the apex. Subsequently, we disrupted both genes individually or concurrently by inserting a selection cassette through homologous recombination, resulting in a loss of corresponding transcripts (**Fig. S4**). No strong difference was visible between single mutants and the wild type, except in the Δ*PpCYP73A49* lines where the phyllids (leaf-like structures) at the apex appeared thinner and more fragile (**Fig. 3F**; **Fig. S5**). In contrast, the Δ*PpCYP73A48/CYP73A49* double mutant consistently exhibited a very strong developmental phenotype characterized by restricted gametophore growth, phyllid fusion and an overproduction of protonema (**Fig. 3F**; **Fig S5**). It can be noted that no major changes in the expression of *PpCYP73A48* and *PpCYP73A49* were observed in single mutant backgrounds (**Fig. S6**). We then attempted to chemically complement the double mutant by providing exogenous *p*-coumaric acid. This treatment successfully restored phyllid expansion compared to mock treatment (**Fig. 3G**; **Fig. S7**), indicating that the Δ*PpCYP73A48/CYP73A49* phenotype was, at least in part, caused by a shortage in *p*-coumaric acid, the product of C4H activity.

### *CYP73* deficiency leads to phenylpropanoid shortage and cuticle defect

Next, we examined the metabolic consequences of C4H deficiency in the moss. UHPLC-UV fingerprints of Δ*PpCYP73* mutant crude extracts revealed the complete loss of UV-absorbing molecules in double mutants (**Fig. 4A**), the major ones being hydroxycinnamate esters as reported before (17, 18). We also noticed a moderate decrease in the main peaks in Δ*PpCYP73A49* mutants as compared to Δ*PpCYP73A48* and the wild-type plants (**Fig. 4A**). These findings were confirmed through targeted UHPLC-MS/MS analysis of diagnostic phenylpropanoids in both crude and saponified extracts (**Fig. 4B-C**). As a result of complete *CYP73* deficiency, none of the targeted molecules downstream of the C4H was detected in metabolic extracts from Δ*PpCYP73A48/CYP73A49* double mutants, whereas we observed concomitant accumulation of *t*-cinnamic acid, the C4H substrate, in saponified extracts (**Fig. 4B-C**). Although no notable changes in metabolic profile of Δ*PpCYP73A48* mutants were observed, a significant decrease in *p*-coumaroyl-5-*O*-shikimate, caffeoyl-2-*O*-threonate and caffeoyl-4-*O*-threonate level was evident in Δ*PpCYP73A49* crude extracts. Corresponding saponified samples showed the accumulation of *t*-cinnamic acid and a corollary reduction in the amount of caffeic acid, while *p*-coumaric acid level remained unchanged (**Fig. 4C**).

**Figure 4.**
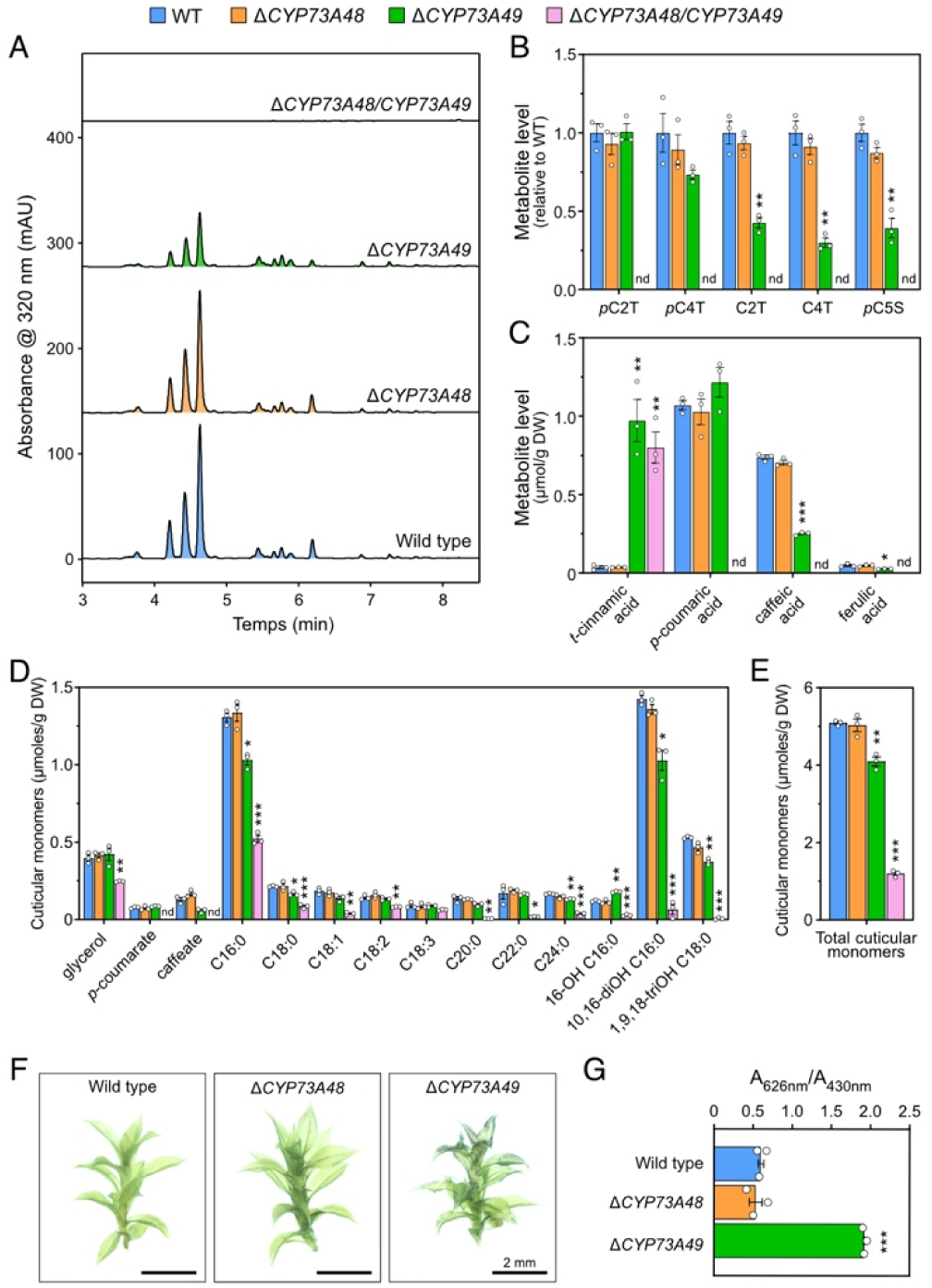
Metabolic and physiological characterization of *P. patens* Δ*CYP73* mutants. (A) Representative UHPLC-UV chromatograms of two-month-old wild-type, Δ*PpCYP73A48*, Δ*PpCYP73A49* and Δ*PpCYP73A48/CYP73A49* crude extracts showing the loss of UV-absorbing molecules in the double mutants. (B) Relative levels of threonate and shikimate phenolic esters in two-month-old gametophore crude extracts. *p*C2T, *p*-coumaroyl-2-*O*-threonate; *p*C4T, *p*-coumaroyl-4-*O*-threonate; C2T, caffeoyl-2-*O*-threonate; C4T, caffeoyl-4-*O*-threonate; *p*C5S, *p*-coumaroyl-5-*O*-shikimate. (C) Quantification of hydroxycinnamic acids in gametophores following the saponification of crude metabolic extracts. (D) Compositional analysis of two-month-old wild-type, Δ*PpCYP73A48*, Δ*PpCYP73A49* and Δ*PpCYP73A48/CYP73A49* gametophore cuticular polymer. (E) Total amount of cuticular monomers from gametophores. (F) Representative pictures of two-month-old wild-type, Δ*PpCYP73A48* and Δ*PpCYP73A49* gametophore after toluidine blue permeability assay. (G) Quantification of toluidine blue permeability. Results in panels B, C, D, E and F are the mean ± SEM of three independent WT biological replicates and three independent mutant lines. WT versus mutant unpaired *t* test adjusted P-value: \**P*<0.05; \*\**P*<0.01; \*\*\**P*<0.001. nd, not detected.

We then examined how impairing C4H function affected the composition of the *P. patens* gametophore cuticular polymer, as its formation relies heavily on phenylpropanoid precursors (17, 18). The analysis of polymer monomers revealed a significant decrease in all aliphatic components, except C18:3, in the Δ*PpCYP73A48/CYP73A49* double mutants compared to wild type (**Fig. 4D**). This decrease was accompanied by a complete loss of the two phenolic monomers, *p*-coumarate and caffeate. The overall quantity of the cuticular polymer, represented by the sum of individual monomers, dropped by 75% in the double mutants compared to the wild-type level (**Fig. 4E**). There were no changes in cuticular polymer composition of Δ*PpCYP73A48* single mutants, consistent with the soluble phenylpropanoid analysis (**Fig. 4A-E**). In contrast, in Δ*PpCYP73A49* single mutants, there were moderate yet statistically significant variations in cuticular polymer composition (**Fig. 4D**), resulting in a *ca*. 20% decrease in total monomer level compared to wild type (**Fig. 4E**). As for phenolic monomers, *p*-coumarate remained unchanged in Δ*PpCYP73A49* mutants but caffeate level was reduced by 50% in the polymer, although this finding lacked statistical support (adjusted P-value = 0.084). To understand the consequences of these polymer composition changes on cuticle properties, we performed toluidine blue assays on gametophores of single Δ*PpCYP73* mutants, as the stunted growth of double Δ*PpCYP73A48/CYP73A49* gametophores prevented their inclusion in the experiment. As shown in **Figure 4F-G**, we observed a significant increase in Δ*PpCYP73A49* gametophore permeability to toluidine blue, indicating a strong defect in cuticle diffusion barrier function despite the moderate compositional changes (**Fig. 4D**). This function appeared unaltered in the Δ*PpCYP73A48* mutant as compared to wild type (**Fig. 4F-G**).

### *CYP73* function is conserved in liverworts and hornworts

To achieve two primary objectives – enhancing our evolutionary conclusions and filling the knowledge gap in bryophytes – we conducted an expanded functional analysis of the *CYP73* family, focusing on the liverwort *M. polymorpha*. We identified the *MpCYP73A1* gene as the primary paralog based on its expression level (**Fig. 5A**-B), for which we also previously confirmed that the corresponding protein catalyzes C4H activity *in vitro* (**Fig. 1D**). We therefore isolated two independent CRISPR mutants of the *MpCYP73A1* gene, using two distinct protospacers located in the first exon. The resulting *Mpcyp73a1-1* and *Mpcyp73a1-2* alleles featured 12 and 37 nucleotide deletions (**Fig. S8**), respectively, leading to four amino acid loss (residues 94-97) and a premature STOP codon (position 150) at the protein level. The impact of these mutations on *M. polymorpha* development was significant, leading to a dramatic reduction in both the thallus area and number of gemmae cups per plant compared to the wild type (**Fig. 5C-E**). Noteworthy, disrupting *MpCYP73A1* function did not affect the expression of the two other *MpCYP73* paralogs, which remained 28- to 243-times lower in the thallus (**Fig. 5B**).

**Figure 5.**
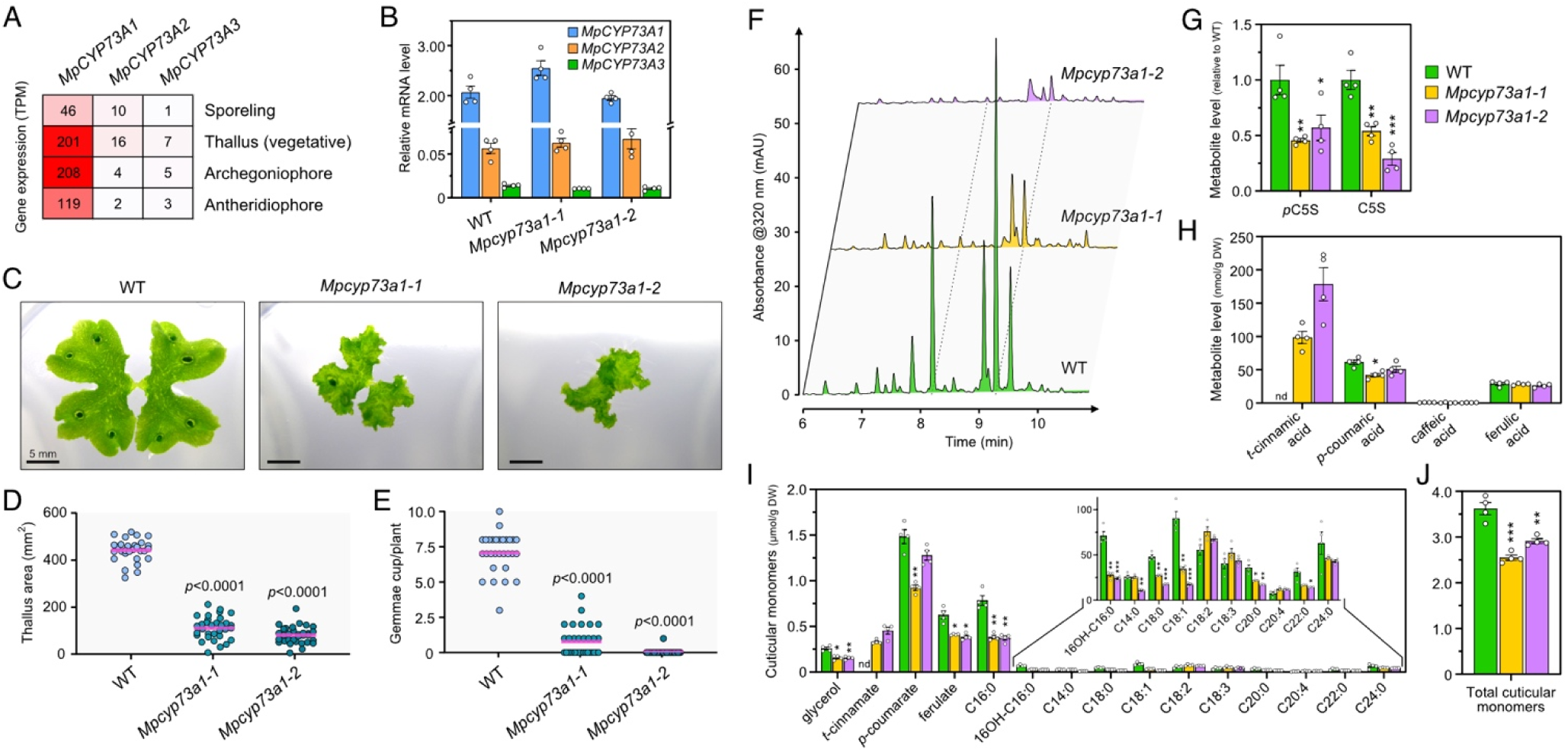
Developmental and metabolic consequences of *CYP73A1* deficiency in *M. polymorpha*. (A) Expression profiles of the three *M. polymorpha CYP73* paralogs in various tissues. Data are derived from the CoNekT database (65). (B) RT-qPCR monitoring of the three *MpCYP73* genes in wild type, *Mpcyp73a1-1* and *Mpcyp73a1-2* mutants. Results are the mean ± SEM of four biological replicates. (C) Representative pictures of three-week-old wild-type, *Mpcyp73a1-1* and *Mpcyp73a1-2* plants. (D) Quantification of three-week-old wild type, *Mpcyp73a1-1* and *Mpcyp73a1-2* thallus area. (E) Quantification of gemmae cup number in three-week-old wild-type, *Mpcyp73a1-1* and *Mpcyp73a1-2* plants. Results are the mean (pink trait) of measurements made on 29-32 plants. Adjusted P-value from WT versus mutant unpaired *t* test is shown. (F) Representative UHPLC-UV chromatograms of one-month-old wild-type, *Mpcyp73a1-1* and *Mpcyp73a1-2* crude extracts. (G) Relative quantification of shikimate phenolic esters in one-month-old thallus crude extracts. C5S, caffeoyl-5-*O*-shikimate; *p*C5S, *p*-coumaroyl-5-*O*-shikimate. (H) Quantification of hydroxycinnamic acids following saponification of crude metabolic extracts. (I) Compositional analysis of one-month-old wild-type, *Mpcyp73a1-1* and *Mpcyp73a1-2* cuticular polymer. (J) Total amount of cuticular monomers. Results in panels G, H I and J are the mean ± SEM of four independent biological replicates for each genotype. WT versus mutant unpaired *t* test adjusted P-value: \**P*<0.05; \*\**P*<0.01; \*\*\**P*<0.001. nd, not detected.

We went on with the metabolic analysis of *Mpcyp73a1* mutants. Through UHPLC-UV fingerprints of crude metabolic extracts, we observed a decrease in main peaks in mutants as compared to wild type (**Fig. 5F**). Targeted UHPLC-MS/MS analysis demonstrated *ca*. 50% reduction in the two essential intermediates of the phenylpropanoid pathway, *p*-coumaroyl-5-*O*-shikimate and caffeoyl-5-*O*-shikimate (18), in *Mpcyp73a1* CRISPR lines (**Fig. 5G**). Furthermore, saponification of the crude extracts, followed by targeted analysis, revealed an accumulation of *t*-cinnamic acid associated with the mutation of *MpCYP73A1* (**Fig. 5H**). However, no significant changes in other hydroxycinnamic acids were observed in the *Mpcyp73a1* mutant lines compared to the wild type, suggesting that the UV-absorbing peaks in *M. polymorpha* crude extracts were likely not related to hydroxycinnamic acid derivatives but rather to other phenylpropanoid classes (e.g., auronidins) (19). We extended the metabolic characterization of *Mpcyp73a1* mutants to the compositional analysis of their cuticular polymer. This analysis uncovered phenolics as the most abundant monomers in *M. polymorpha* (**Fig. 5I**). Both mutants consistently accumulated *t*-cinnamate in their polymer and featured a *ca*. 35% reduction in ferulate compared to the wild type. Impairing *MpCYP73A1* function also caused alterations in the aliphatic monomer composition of the cuticular polymer, including decrease in glycerol, palmitate (C16:0), and ω-hydroxypalmitate (16OH-C16:0) (**Fig. 5I**). Overall, the cuticular polymer load was significantly reduced in *Mpcyp73a1* mutant lines, with a reduction of up to 25% (**Fig. 5J**).

In the final stage of our study, we performed pharmacological experiments using the selective inhibitor piperonylic acid (PA) to disrupt C4H in planta (20, 21). PA treatment accurately replicated in *P. patens* wild-type plants the developmental phenotype observed in Δ*PpCYP73A48/CYP73A49* double mutants, displaying stunted gametophores and protonema outgrowth (**Fig. 6A**). Similarly, we observed a dramatic inhibition by PA of *M. polymorpha* thallus expansion, which resembled an aggravation of *Mpcyp73a1* single mutant phenotypes (**Fig. 6A**). Based on aforementioned findings, we reasoned that PA treatment could serve as a surrogate for gene inactivation and therefore applied the pharmacological approach to investigate C4H function in hornworts, the third bryophyte group. To explore this, we treated the hornwort *Anthoceros agrestis* with PA, considering the existence of biochemical evidence of CYP73-catalyzed C4H activity in this species (22). The PA treatment triggered a detectable change in *A. agrestis* thallus development, characterized by inhibition of expansion and the development of rounded and thickened edges (**Fig. 6A**). Moreover, PA treatment caused an extensive flattening of UHPLC-UV chromatograms compared to mock treatment (**Fig. 6B**), suggesting that phenylpropanoid biosynthesis was impaired. This finding was further confirmed by UHPLC-MS/MS targeted analysis of crude extracts, which revealed the almost complete loss of *p*-coumaroyl-5-*O*-shikimate and caffeoyl-5-*O*-shikimate, as well as a *ca*. 50% decrease in rosmarinic acid, a caffeate ester (23), upon PA treatment (**Fig. 6C**). Analysis of saponified extracts underscored the accumulation of *t*-cinnamic acid and significant reduction in other hydroxycinnamic acids downstream of the C4H step in PA-treated plants compared to the mock treatment (**Fig. 6D**). Since *cis*-cinnamic, a *t*-cinnamic acid stereoisomer, was suggested to play a role in the developmental abnormalities linked to C4H deficiency in Arabidopsis (24), we conducted a targeted analysis of these two compounds in crude extracts of C4H-impaired plants. As illustrated in **Fig. S9**, while *t*-cinnamic acid was occasionally found to accumulate, *c*-cinnamic acid was not detected in examined samples.

**Figure 6.**
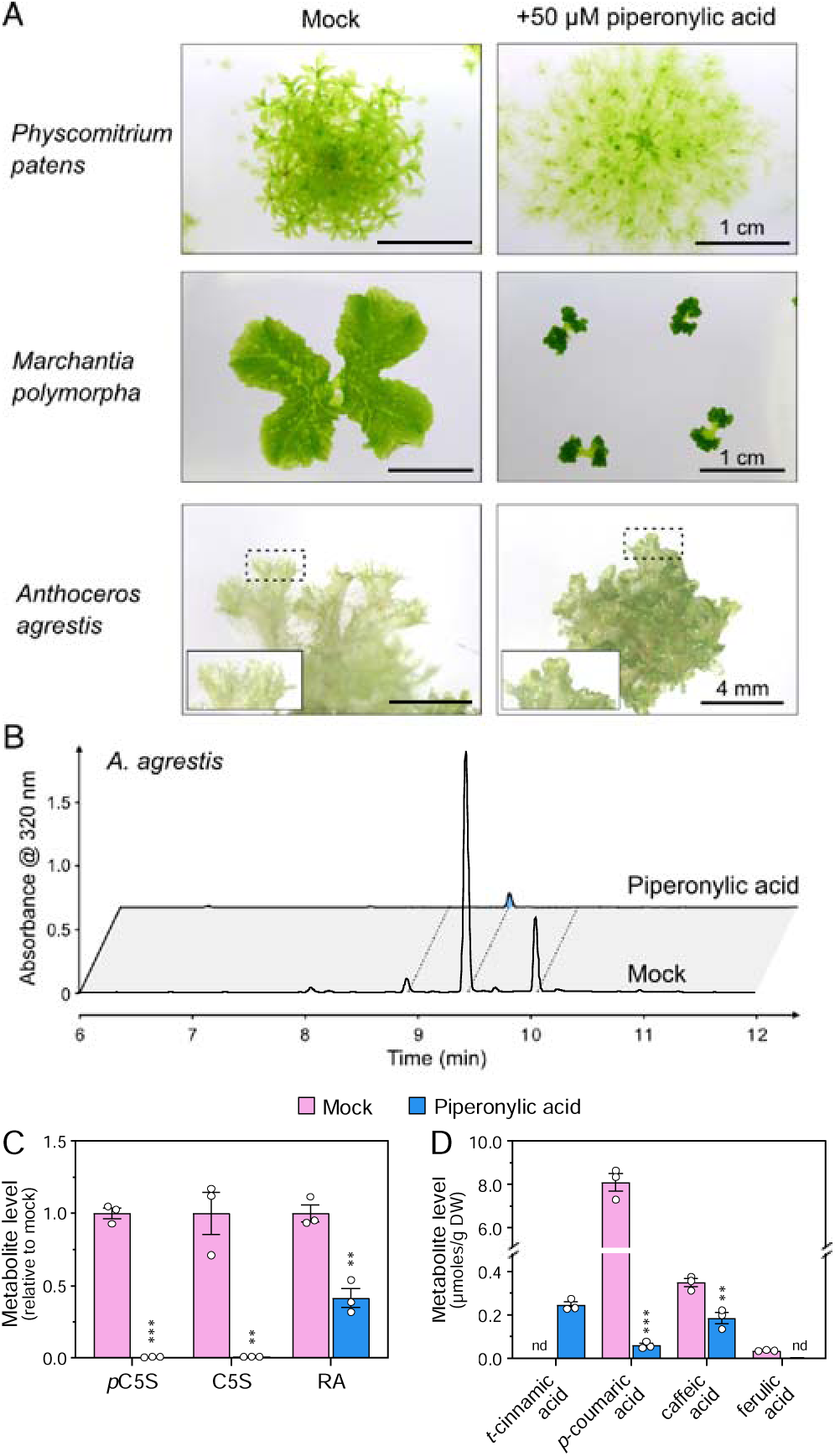
Developmental and metabolic effects of piperonylic acid in bryophytes. (A) Pictures of one-month-old *P. patens*, three-week-old *Marchantia polymorpha* and two-month-old *Anthoceros agrestis* plants treated with 50 µM piperonylic acid (PA) for the entire culture period. Mock treatment was performed with 0.05% DMSO. (B) Representative UHPLC-UV chromatograms of mock- and PA-treated *A. agrestis* metabolic extracts. (C) Relative quantification of phenolic esters in mock- and PA-treated *A. agrestis* plants. C5S, caffeoyl-5-*O*-shikimate; *p*C5S, *p*-coumaroyl-5-*O*-shikimate; RA, rosmarinic acid. (D) Quantification of hydroxycinnamic acids *in A. agrestis* plants following saponification of metabolic extracts. Results in panels C and D are the mean ± SEM of three independent biological replicates for each condition. Mock versus PA unpaired *t* test adjusted P-value: \*\**P*<0.01; \*\*\**P*<0.001. nd, not detected.

## DISCUSSION

The phenylpropanoid pathway is an essential plant metabolism unequivocally linked to embryophyte evolution in its canonical form. It contributes both soluble molecules and precursors of structural, insoluble polymers that address the challenges posed by terrestrial life. It is estimated that tree species can direct up to 30% of photosynthetic carbon into this pathway, as inferred from lignin proportion in wood (25, 26), stressing the pivotal role of the phenylpropanoid pathway in embryophyte biology and Earth biogeochemical cycles as carbon sink. Such a substantial flux certainly necessitates a tight metabolic organization and regulation. In this context, the first three steps assume critical importance as they essentially handle the entirety of this flux. While the implication of the phenylpropanoid pathway in plant adaptation to terrestrial constraints is well-documented in tracheophytes, very little was known about its origin and its function in bryophytes, the non-vascular embryophytes. Here we report the evolution reconstruction and functional analysis in bryophyte species of the *CYP73* gene family that encodes the second step of the phenylpropanoid pathway, the *t*-cinnamic acid 4-hydroxylase (C4H).

Based on both computational and experimental data, our updated evolutionary investigation in a broad set of plant genomes provides evidence that C4H activity emerged in an embryophyte progenitor together with the *CYP73* gene family. These findings resonate with previous studies showing that potential *CYP73* orthologs are exclusive to embryophytes (10, 27, 28). Whether this evolutionary scenario will stand valid in light of upcoming, new genomic data from algae remains a question. Nonetheless, the evolutionary pattern of the *CYP73* family diverges from that of other genes within the phenylpropanoid pathway, ranging from *PAL* to *CYP98*, for which distant homology is evident in charophytes (27–29). The seemingly sudden and delayed emergence of the *CYP73* gene family thus raises questions about the evolutionary path and mechanisms that gave birth to this gene family. It also suggests that *CYP73* emergence was instrumental in founding the canonical phenylpropanoid pathway in an embryophyte ancestor, in line with the often observed contribution of new CYP families to creating novel metabolic pathways (13, 30). Moreover, previous molecular evolution analysis underscored the consistent trend of strong purifying selection acting on *CYP73* genes across both tracheophytes and bryophytes, hinting at the fact they fulfilled essential function from the early stages of land plant evolution (10).

Together with data from the study of the Arabidopsis *CYP73A5* gene (14), our functional analysis of bryophyte *CYP73* genes establishes the C4H step as a critical and evolutionarily conserved element of the canonical phenylpropanoid pathway. The recruitment and fixation of C4H within embryophytes delineates an intriguing facet, particularly when considering the potential of some plant enzymes to establish an alternative route to *p*-coumaric acid. A prominent example are the bifunctional phenylalanine/tyrosine ammonia-lyases (PTAL), which derive from to the ubiquitous PAL enzyme family. In the monocot *Brachypodium distachyon*, PTAL bypasses C4H and contributes to nearly half of lignin production through L-tyrosine (31). However, PTAL are mainly restricted to grasses, and *B. distachyon* retained three *CYP73* genes that encode functional C4H enzymes (10). The precise benefits of positioning and conserving C4H as a central module in the phenylpropanoid pathway is elusive. Exploration of catalytic properties revealed the high affinity (*K*_m_ < 10 µM) and relatively slow turnover rates of C4H enzymes across various species (10, 32–34). Furthermore, it was observed that C4H substrate preference is constrained, being capable of using only *t*-cinnamic acid mimics as substrates (20, 33, 35, 36). Thus, the catalytic attributes of C4H potentially prevent metabolic derailment that could result from enzyme promiscuity, while concurrently offering the potential for efficient control over metabolic flux. This perspective is further supported by the documented role of the ER membrane-bound C4H as a nucleation point for a phenylpropanoid metabolon (8, 37), facilitating the recruitment of other CYP and soluble enzymes within close proximity for the effective and dynamic channeling of metabolic intermediates.

Our work demonstrates that perturbing C4H function, either via *CYP73* gene inactivation or inhibitor treatment, has a significant impact on bryophyte development. This aligns with the findings obtained in the tracheophyte Arabidopsis, where interfering with the five first steps of the phenylpropanoid pathway consistently led to lignin alteration and growth reduction (14, 38–41). Four hypotheses were proposed to explain this phenomenon, ranging from impaired water transport due to xylem collapse, negative feedback through a cell wall surveillance system, and over- or under-accumulation of phenylpropanoid derivatives (reviewed in (42)). Whether the developmental alterations we observed in bryophytes, which lack lignified vascular tissues, falls under some of these hypotheses is an interesting question. It’s worth noting that the impact of impeding C4H function in bryophytes extends beyond mere growth reduction. It profoundly alters the developmental plan, leading for instance in *P. patens* to phyllid expansion failure and organ fusion symptoms, as observed in mutants of downstream steps (17, 18). Interestingly, this strong developmental phenotype could be mitigated by supplying exogenous *p*-coumaric acid. This suggests that the observed phenotypes were unlikely due to hyperaccumulation of *t*-cinnamic acid derivatives. As proposed for other *P. patens* phenylpropanoid mutants (17, 18), phenylpropanoid shortage might account for the developmental anomalies in C4H-impaired bryophytes. This hypothesis could be linked to cuticle defects, which might function, in absence of lignin and suberin, as a structural element in bryophytes, contributing to tissue scaffolding. This perspective does not necessarily contradict the two remaining hypotheses – feedback control through cell wall monitoring system or impaired water transport. However, the latter hypothesis appears less substantiated, as neither growing *P. patens* Δ*PpCYP73* mutants in liquid culture nor embedding them in agarose medium could rescue their stunted phenotype. Further investigations will be necessary to elucidate the factors driving the developmental consequences of C4H deficiency in bryophytes and to determine whether these mechanisms are conserved across embryophytes.

## MATERIALS AND METHODS

### Ortholog search and phylogenetic reconstruction

The investigation of *CYP73* orthology involved a reciprocal best hits (RBH) method, where the AtCYP73A5 protein sequence was used as the initial query (**see Tab. S1 for RBH output**). Briefly, we conducted a forward BLASTp search across 20 Viridiplantae genomes. Next, for each species, the top hit based on the bit-score was used to perform a reverse BLASTp search against the *Arabidopsis thaliana* genome. The presence of a potential *AtCYP73A5* ortholog was considered positive when AtCYP73A5 was identified as the best hit in the reverse search. To update our previously published *CYP73* phylogeny (10), we incorporated 72 new sequences retrieved from recently released genomes or through an extended search in the 1kP transcriptome dataset (43), resulting in a total of 275 sequences. Alignment of the sequences was performed using the MUSCLE algorithm (44) based on their protein sequences (**full alignment available as supplemental Data S1**). Ambiguous regions in the obtained alignment were masked using the Gblocks (45) implemented via Seaview (46), allowing less strict flanking positions. The *CYP73* phylogeny was then reconstructed from the Gblocks nucleotide alignment using IQ-TREE2 v2.2.2.6 (47). The reconstruction employed the following command line: “./iqtree2 -s Gblock_alignment.fst --alrt 1000 -B 1000”, which implements ModelFinder for model selection and computes ultrafast bootstrap and SH-aLRT tests with 1,000 pseudo-replicates to determine branch support. The GTR+F+I+R7 model was used for phylogenetic reconstruction (**tree file available as supplemental Data S2**).

### Homology modelling and docking experiments

Tridimensional structure of transmembrane segment-free PpCYP73A48 protein (residues 46-517) was built using the SWISS-MODEL server homology modelling pipeline (48). Both BLAST and HHblits were used to search for the best template in SWISS-MODEL library. Models were built based on the target-template alignment using ProMod3 v3.1.1 and *Sorghum bicolor* C4H1 structure (pdb: 6VBY) as template. The resulting structure of PpCYP73A48 exhibited a good structural alignment with SbC4H1, with a global QMEANDisCo score of 0.87 ±0.05. Heme prosthetic group was first docked into reconstructed PpCYP73A48 3D model using Autodock Vina (49) in flexible mode, allowing geometry of cysteine 459 to change. Resulting heme-containing PpCYP73A48 3D model (**pdb file available as supplemental Data S3**) was subsequently used for *t*-cinnamic acid docking via Autodock Vina in rigid mode. For *t-*cinnamic acid, a 40 Å × 40 Å × 40 Å search box positioned above the heme was used. Ligand and receptor files required for docking experiments were prepared with AutodockTools 1.5.7 (https://ccsb.scripps.edu/mgltools/). Docking results were visualized with ChimeraX software (50).

### Generation of yeast expression plasmids

Coding sequences (CDS) of *Physcomitrium patens CYP73A48*, *CYP73A49* and *CYP73A51* were PCR-amplified from *P. patens* Gransden cDNA and cloned into the yeast expression plasmid pYeDP60 by restriction/ligation using *Bam*HI/*Kpn*I (CYP73A48 and CYP73A51) or *Sma*I/*Kpn*I (CYP73A49) restriction sites. *Marchantia polymorpha CYP73A1* coding sequence was PCR-amplified from *M. polymorpha* Tak-1 cDNA using Gateway-compatible primers. Yeast codon-optimized *Phaeoceros carolinianus CYP73A* (*PcCYP73A*; RXRQ_scaffold_2140006) and native *Klebsormidium nitens kfl00038_0230* coding sequences were synthesized as double-strand DNA fragments (gBlock, Integrated DNA Technologies) containing Gateway *att*B1 and *att*B2 extensions (**sequences available in Tab. S2**). After generation of pENTRY plasmids by BP recombination, *MpCYP73A1*, *PcCYP73A* and *kfl00038_0230* CDS were shuttled to a Gateway version of pYeDP60 by LR recombination. For 3xHA tagging of CYP73 proteins, STOP codon-free CDS were PCR amplified using Gateway-compatible primers and cloned into pDONR207 by BP recombination. CDS were transferred to a modified pAG425GAL-ccdB-3xHA yeast expression plasmid (51), in which the *LEU2* auxotrophic marker was replaced by *ADE2*. Plasmids containing yeast-optimized CDS of *Brachypodium distachyon CYP73A92* and *CYP73A94* were described previously (10). For expression of CYP73 mutated proteins, site-directed mutagenesis was performed on pYeDP60 plasmids by overlapping PCR, using primers containing the desired mutations. To reduce background originating from methylated template plasmids, PCR reactions were treated with *Dpn*I prior to *E. coli* transformation.

### Production of recombinant CYP73 proteins in yeast

Yeast expression plasmids harboring *CYP73* CDS were introduced into the *Saccharomyces cerevisiae* WAT11 strain (52). Procedures for recombinant CYP expression and microsome purification were the same as in (12). Total microsomal protein concentration was determined according to the Qubit™ protein assay kit (ThermoFisher Scientific). Recombinant CYP proteins were quantified by type II differential spectrophotometry using reduced cytochrome P450 and carbon monoxide (53). Expression of CYP73-3xHA tagged proteins was analyzed by Western blot. To this end, microsomal proteins were denatured for 5 min at 65°C in Laemmli buffer containing DTT as reducing agent. Proteins were quantified by the amido black protein assay with bovine serum albumin as reference. Ten micrograms of denatured microsomal proteins were separated by SDS-PAGE using 10% polyacrylamide gel (Mini-PROTEAN TGX precast gel, Biorad) and tris/glycine buffer. After electrophoresis, proteins were transferred on Immobilon-P PVDF membrane. HA-tagged proteins were detected using primary antibody anti-HA (1/10 000) followed by incubation with the HRP-conjugated secondary antibody and finally revealed by chemiluminescence (Clarity Western ECL, Biorad) on a Fusion-FX imager (Vilber).

### *In vitro* enzyme assays

Standard assay for cinnamic acid 4-hydroxylase activity was performed in a 100 µL reaction containing 50 mM potassium phosphate buffer (pH 7.4), 150 µg microsomal proteins (∼5 µl microsomal preparations), 100 µM *trans*-cinnamic acid (from a 10 mM stock solution prepared in DMSO) and 500 µM NADPH. The reaction was initiated by addition of NADPH and incubated at 28°C in the dark for 30 min. Reaction was stopped by the addition of 100 µl methanol followed by thorough agitation. Samples were centrifuged for 10 min at 15,000*g*, 4°C to pellet microsomes. Supernatant was recovered and used for UHPLC-MS/MS analysis. For *in vitro* assay with mutated CYP73 enzymes, same procedures were followed except that 5 pmoles P450 enzyme per assay were used, and samples were analyzed by HPLC-UV. The *t*-cinnamic acid binding properties of CYP73 recombinant proteins were assayed spectrophotometrically by monitoring the low-spin to high-spin transition upon substrate binding in the active site (i.e., type I spectra). To this end, absorbance difference in the 380-500 nm range was measured with 150 nM recombinant CYP73 proteins and 100 µM *t*-cinnamic acid.

### Plant material and growth conditions

In this study we used Physcomitrella (*Physcomitrium patens*) Gransden accession (54), *Anthoceros agrestis* Bonn isolate (55) and *Marchantia polymorpha* Tak-1 accession (56). The Arabidopsis *cyp73a5-1* mutant was previously characterized (10). *P. patens* and *A. agrestis* were grown axenically in Knop medium as described before (17). *M. polymorpha* was grown axenically on half-strength Gamborg B5 medium (Duchefa, #G0209) containing 0.5 g/L MES as pH buffering agent. Gamborg B5 medium was adjusted to pH 5.5 with KOH and solidified with 12 g/L agar (#4807, Roth). Plants were grown in 93 × 21 mm petri plates filled with 30 mL of medium. Alternatively, *P. patens* was cultivated in 500 mL Erlenmeyer flasks filled with 200 mL liquid Knop medium and sealed with C-type Silicosen® stoppers. *P. patens* liquid cultures were constantly agitated at 130 rpm on an orbital shaker. *Arabidopsis* plants were grown on soil in 7 cm square pots. Plants were kept under a 22/18°C, 16h/8h light/dark regime. Bryophyte species and Arabidopsis were exposed to 50 µmol/m^2^/s and 100 µmol/m^2^/s light intensity, respectively, provided by 20W/840 white cool LED tubes (Philips).

### Generation of *P. patens* transgenic lines

Δ*PpCYP73s* knock-out mutants were generated via homologous recombination-mediated gene disruption as described previously (18). *CYP73A48* and *CYP73A49* disruption constructs were excised from vector backbone by *Bam*HI and *Kpn*I digestion, respectively. The Δ*PpCYP73A48/CYP73A49* double mutants were produced by disrupting the *PpCYP73A49* gene in the Δ*PpCYP73A48* #23 mutant background. For *uidA* reporter lines, two genomic regions for homologous recombination framing the STOP codon were PCR-amplified from genomic DNA and assembled with the *uidA* reporter gene following the same procedures as in (18). The *CYP73A48:uidA* and *CYP73A49:uidA* constructs were excised from vector backbone by *Nhe*I digestion. 25 µg of excised fragment were used for protoplasts transfection. Since *CYP73:uidA* constructs do not contain a selection marker, it was co-transfected with the pRT101 plasmid containing the *NPTII* selection cassette (57). Transformants were selected on Knop plates supplemented with 25 mg/L geneticin (Δ*PpCYP73A48* mutants and *PpCYP73A:uidA lines*) or 10 mg/L Hygromycin B (Δ*PpCYP73A49* mutants). Transgenic lines were molecularly characterized as described previously (18).

### Arabidopsis *cyp73a5-1* mutant trans-complementation

Coding sequences of *CYP73A48*, *CYP73A49*, *CYP73A51* and *CYP73A5* were transferred by LR recombination into the Gateway pCC1061 binary vector, which contains a 2977 bp promoter fragment from Arabidopsis *CYP73A5* gene (58). Obtained expression plasmids were introduced into *Agrobacterium tumefaciens* C58c1 strain and used to transform heterozygous *cyp73a5-1* plants by the floral dip method (59). Transformants were selected based on their resistance to kanamycin. T1 plants homozygous for the *cyp73a5-1* allele were identified by PCR (**primers in Tab. S2**) and further confirmed by the full resistance of T2 progeny to sulfadiazine. Experiments were performed with T3 plants homozygous for both the mutant allele and the trans-complementation construct.

### GUS staining

Plant tissues were vacuum infiltrated during 10 min with a GUS solution containing 50 mM potassium phosphate buffer pH 7.0, 0.5 mM ferrocyanide, 0.5 mM ferricyanide, 0.1% Triton X-100 and 0.5 mg/mL X-Gluc, and incubated at 37°C for 4.5 h. Chlorophyll was removed by washing tissues three times in 70% ethanol.

### Chemical complementation with *p*-coumaric acid

Protoplasts from *P. patens* wild type and Δ*PpCYP73A48/CYP73A49* mutant were embedded in low-melting point agarose as described before (60). After three days, regeneration solution that overlaid the solidified film was changed to Knop medium supplemented 50 µM *p*-coumaric acid. Mock treatment was performed by supplementing Knop medium 0.1% ethanol. Regeneration of protoplasts was performed in standard growth conditions and monitored over five weeks.

### Soluble metabolite extraction and analysis

Tissue collection, lyophilization and grinding were performed as described before (18). Metabolites were extracted from lyophilized plant material using a methanol:chloroform:water protocol as described previously (18). Briefly, 500 µl methanol were added to 10 mg lyophilized plant material. Samples were agitated for 1h at 1500 rpm at room temperature prior to addition of 250 µl chloroform. After agitation for 5 min, phase separation was induced by addition of 500 µl water followed by vigorous agitation and centrifugation (15000*g*, 4°C, 15 min). Supernatants were recovered and constituted the crude metabolic extracts. To release hydroxycinnamic acid (HCAA) aglycones from corresponding soluble esters, crude metabolic extracts were saponified. To this end, 200 µl of extract were dried in vacuo, resuspended in 200 µl 1M NaOH and incubated for 2h at 30°C under 1000 rpm agitation. NaOH was neutralized with 33.3 µl 6M HCl and HCAA were extracted twice with 1 mL ethyl acetate. Pooled organic phases were dried in vacuo and resuspended in 200 µl 50% MetOH prior to analysis. Unless otherwise stated, metabolites were separated and detected on a Dionex UltiMate 3000 UHPLC (ThermoFisher Scientific) system coupled to an EvoQ Elite LC-TQ (Bruker) mass spectrometer as reported before (18). Molecules were ionized in positive or negative mode via a heated electrospray ionization source (HESI, Bruker) and detected by specific multiple reaction monitoring (MRM) methods (**Tab. S3**). Concurrently to MS/MS data acquisition, UV-absorbance profiles were recorded with a Ultimate 3000 photodiode array detector operated in the 200-400 nm range (ThermoFisher Scientific). UV data were analyzed with Bruker Data Analysis software. In the case of mutated CYP73 enzyme assays, metabolites were analyzed by HPLC-UV on a Alliance 2695 chromatographic system (Waters) coupled to a photodiode array detector (PDA 2996; Waters) as described before (18).

### Determination of cuticular polymer composition

*P. patens* cuticular polymer composition, including glycerol, was determined from two-month-old, lyophilized gametophores grown in liquid culture as previously reported (61). For *M. polymorpha*, same procedures were followed except that heptadecanoic acid (C17:0) and ribitol were used as internal standards. *M. polymorpha* samples were analyzed using gas chromatography (GC; Agilent GC8890) coupled with a time-of-flight mass spectrometer (TOFMS; Leco Pegasus BT2). One microliter of derivatized monomers was injected on a HP-5MS Ultra Inert column (30 m × 0.25 mm × 0.25 µm; Agilent) in 100:1 split mode. The temperature gradient was set as follows: 120°C for 1 min, 120°C to 340°C at 10°C/min, and 340°C for 3 min. Helium was used as the carrier gas at a flow rate of 1 mL/min. Transfer line and source temperatures were maintained at 250°C. Analytes were ionized and fragmented by electronic impact at 70 eV. Data were acquired over a *m/z* 45-500 mass range with a frequency of 8 spectra/s. Compounds were identified according to two orthogonal criteria: spectral similarity score (>800) and retention index (±20). After identification and peak area integration, compound quantification was carried out using the internal standards (ribitol, 5TMS and C17:0, 1TMS) and response factors obtained from the analysis of authentic standards. When standards were unavailable, response factors from structurally-related molecules were utilized (**see Tab. S4**).

### Tissue permeability assay

*P. patens* tissue permeability was probed by immersing gametophores for 30s in a 0.05% toluidine blue solution containing 2% tween20. Care was taken to remove rhizoids prior to staining assay and not to immerse the cut area. Gametophores were then abundantly rinsed with distilled water. For each genotype or replicate, ten gametophores were individually processed and subsequently pooled. Sample pigments were extracted in 400 µl of buffer (200 mM Tris-HCl, 250 mM NaCl, 25 mM EDTA) with two 3 mm steel balls operated at 30 Hz for 5 min. Following addition of 800 µl ethanol, samples were vortexed and centrifuged for 15 min at 18000*g*. Absorbance of supernatants at 626 nm (A626) and 430 nm (A430) were recorded; A626/A430 ratio was used to quantify toluidine blue levels in plant tissue.

### Gene expression analysis by RT-qPCR

Total RNA was extracted from five-week-old, lyophilized *P. patens* gametophores and *M. polymorpha* thalli using TriReagent (Sigma) and was subsequently treated with RQ1 DNAse (Promega). 150 ng (*P. patens*) or 1 µg (*M. polymorpha*) of DNase-treated total RNA was retro-transcribed with the SuperScript IV enzyme (ThermoFisher Scientific) and an oligo(dT)18, following the manufacturers’ instructions. qPCR was performed in a 384-well plate, in a reaction volume of 10 µL containing 1 µl (*P. patens*) or 0.2 µl (*M. polymorpha*) RT reaction, 0.5 µM of each primer, and 5 µL of Master mix SYBR Green I (Roche). Reactions were run in triplicates using a LightCycler^®^ 480 II (Roche) with the following program: 95°C for 10 min, followed by 40 amplification cycles [95°C for 10 s (denaturation) – 60°C for 15 s (hybridization) – 72°C for 15 s (extension)], and a melting curve analysis from 55 to 95°C to verify primer specificity. Crossing points (Cp) were determined using manufacturer’s software and corrected with primer pair efficiency. The expression levels of *P. patens CYP73* genes were normalized to the expression levels of *PpTRX1* (*Pp3c19_1800*) and *PpSEC15* (*Pp3c27_3270*); the expression levels of *M. polymorpha CYP73* genes were normalized to the expression levels of *MpACT7* (*Mp6g11010*) and *MpEF1* (*Mp3g2340*).

### *M. polymorpha* CRISPR/Cas9-mediated genome editing

Mutant alleles of *M. polymorpha CYP73A1* were generated by CRISPR/Cas9 following the procedures described in (62). Briefly, two independent protospacer sequences in *CYP73A1* first exon were identified using the CRISPOR tool (63). Selected protospacers started with a G for proper U6 promoter-driven expression and had a specificity score of 100. Double-stranded protospacer fragments were reconstituted by hybridization of two complementary oligonucleotides and were *Bsa*I-cloned into the Gateway pMpGE-En03 vector that contains Mp*U6-1* promoter and gRNA scaffold (62). MpU6pro-gRNA expression cassettes were subsequently transferred by LR recombination into the binary pMpGE011 vector, which contains a *SpCas9* expression cassette and allows chlorsulfuron-based selection of plant transformants (62). Recombined pMpGE011 vectors were introduced into *A. tumefaciens* GV3101 strain and served for the transformation of *M. polymorpha* Tak-1 accession according to the thallus method (64). Transformants were selected on half-strength Gamborg B5 medium supplemented with 0.5 µM chlorsulfuron. Genome editing was checked in transformants by Sanger sequencing of PCR amplicons, using primers designed by the CRISPOR tool. Edited lines were further propagated from a single gemma to avoid mosaicism and confirm mutant allele.

### Piperonylic acid treatment

Standard agar growth media were supplemented with 50 µM piperonylic acid (PA) using 100 mM stock solution prepared in DMSO. Mock medium was prepared by supplementing growth media with 0.05% DMSO. PA treatment was initiated by transferring wild-type *P. patens* gametophores, *M. polymorpha* gemma and *A. agrestis* thallus pieces in PA and mock media.

## ACCESSION NUMBERS

*AtCYP73A5*, *At2g30490*; *BdCYP73A92*, *Bradi2g31510*; *BdCYP73A94*, *Bradi3g43160*; *MpCYP73A1*, *Mp6g00020*; *MpCYP73A2*, *Mp8g05070*; *MpCYP73A3*, *Mp8g05080*; *PpCYP73A48*, *Pp3c4_21680*; *PpCYP73A49*, *Pp3c25_10190*; *PpCYP73A50*, *Pp3c13_14870*; *PpCYP73A51*, *Pp3c3_17840*.

## AUTHOR CONTRIBUTIONS

S.K. and H.R. designed the project. S.K., L.K., K.T., L.M., G.W., B.B. and H.R. performed experiments. S.K., L.K., K.T., B.B. and H.R. analyzed data. T.K. and R.R. provided critical material and protocols. H.R. wrote the manuscript with inputs from all authors.

## COMPETING INTEREST STATEMENT

Authors declare no competing interests.

## ACKNOWLEDGMENTS

HR received support from the initiative of excellence IDEX Unistra (ANR-10-IDEX-0002-02) and the Agence Nationale de la Recherche (ANR-19-CE20-0017). HR is grateful to the “Région Grand Est - Fonds Régional de Coopération pour la Recherche 2019” (VitEst project) for partly funding the GC-TOFMS equipment. RR acknowledges support from the German Research Foundation DFG under Germany’s Excellence Strategy (EXC-2189, CIBSS). HR and RR acknowledge the support of the Freiburg Institute of Advanced Studies FRIAS and the University of Strasbourg Institute of Advanced Study USIAS to the METABEVO project.

## SUPPORTING INFORMATION

**Table S1.** Search for *CYP73* orthologs in *Viridiplantae* genomes by reciprocal best hits (RBH). Forward BLASTp search across 20 *Viridiplantae* genomes was performed using *A. thaliana* CYP73A5 protein sequence as query. For each species, the top hit based on the bit-score was used to perform a reverse BLASTp search against the *Arabidopsis thaliana* genome.

| Plant species | Forward BLASTp |  |  |  | Reverse BLASTp |  |
| --- | --- | --- | --- | --- | --- | --- |
|  | Best hit | Bit-score | e-value | identity | Best hit | Bit-score |
| *Arabidopsis thaliana* | AT2G30490 | 1037 | 0 | 100% | AT2G30490 \| CYP73A5 | 1037 |
| *Brachypodium distachyon* | Bradi2g53470 | 862 | 0 | 80% | AT2G30490 \| CYP73A5 | 839 |
| *Amborella trichopoda* | scaffold00077.91 | 873 | 0 | 83% | AT2G30490 \| CYP73A5 | 853 |
| *Picea abies* | MA_118702g0010 | 814 | 0 | 79% | AT2G30490 \| CYP73A5 | 813 |
| *Thuja plicata* | Thupl.29378841s0014 | 810 | 0 | 79% | AT2G30490 \| CYP73A5 | 814 |
| *Salvinia cucullata* | s0039.g012203 | 755 | 0 | 76% | AT2G30490 \| CYP73A5 | 749 |
| *Ceratopteris richardii* | Ceric.38G063100 | 757 | 0 | 72% | AT2G30490 \| CYP73A5 | 738 |
| *Diphasiastrum complanatum* | Dicom.04G054900 | 782 | 0 | 76% | AT2G30490 \| CYP73A5 | 764 |
| *Selaginella moellendorffii* | 175973 | 714 | 0 | 71% | AT2G30490 \| CYP73A5 | 720 |
| *Physcomitrium patens* | Pp3c4_21680 | 767 | 0 | 73% | AT2G30490 \| CYP73A5 | 747 |
| *Marchantia polymorpha* | Mp6g00020 | 706 | 0 | 71% | AT2G30490 \| CYP73A5 | 718 |
| *Anthoceros agrestis* | 228.5028.1 | 748 | 0 | 72% | AT2G30490 \| CYP73A5 | 743 |
| *Penium margaritaceum* | pm004896g0030 | 116 | 3e-26 | 26% | AT2G26170 \| CYP711A1 | 208 |
| *Zygnema circumcaritanum* | Zci_08574.1 | 162 | 4e-42 | 28% | AT2G40890 \| CYP98A3 | 310 |
| *Mesotaenium endlicherianum* | ME000292S04845 | 115 | 8e-26 | 27% | AT1G31800 \| CYP97A3 | 695 |
| *Spirogloea muscicola* | SM000094S24703 | 111 | 2e-24 | 24% | AT1G31800 \| CYP97A3 | 709 |
| *Chara braunii* | CHBRA18g00060 | 163 | 2e-41 | 35% | AT2G45560 \| CYP76C1 | 187 |
| *Klebsormidium nitens* | kfl00038_0230 | 235 | 1e-69 | 31% | AT2G40890 \| CYP98A3 | 260 |
| *Chlamydomonas reinhardtii* | Cre02.g142266 | 119 | 4e-27 | 25% | AT1G31800 \| CYP97A3 | 590 |
| *Ostreococcus lucimarinus* | gwEuk.21.62.1 | 114 | 7e-26 | 25% | AT1G31800 \| CYP97A3 | 565 |

**Figure S1.**
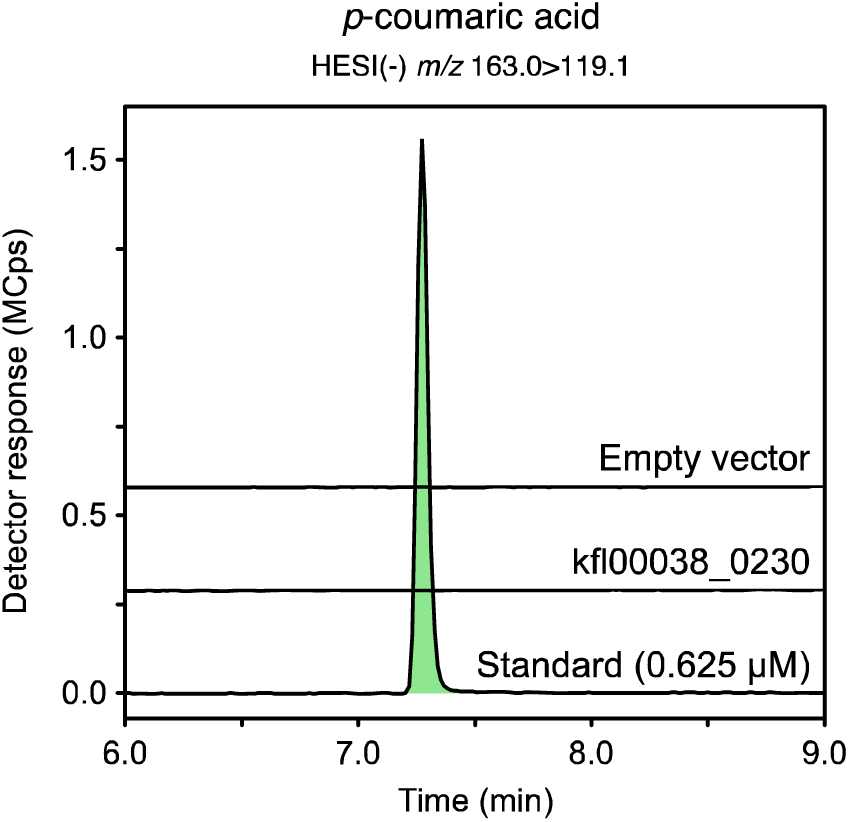
*In vitro* C4H assay with *Klebsormidium nitens* kfl00038_0230 recombinant protein. Representative UHPLC-MS/MS chromatograms showing the absence of *p*-coumaric acid production from *t*-cinnamic acid (C4H activity) when using microsomes from yeasts transformed with the pYeDP60:kfl00038_0230 plasmid. As a negative control, assays were also performed with microsomes from yeasts transformed with an empty vector.

**Figure S2.**
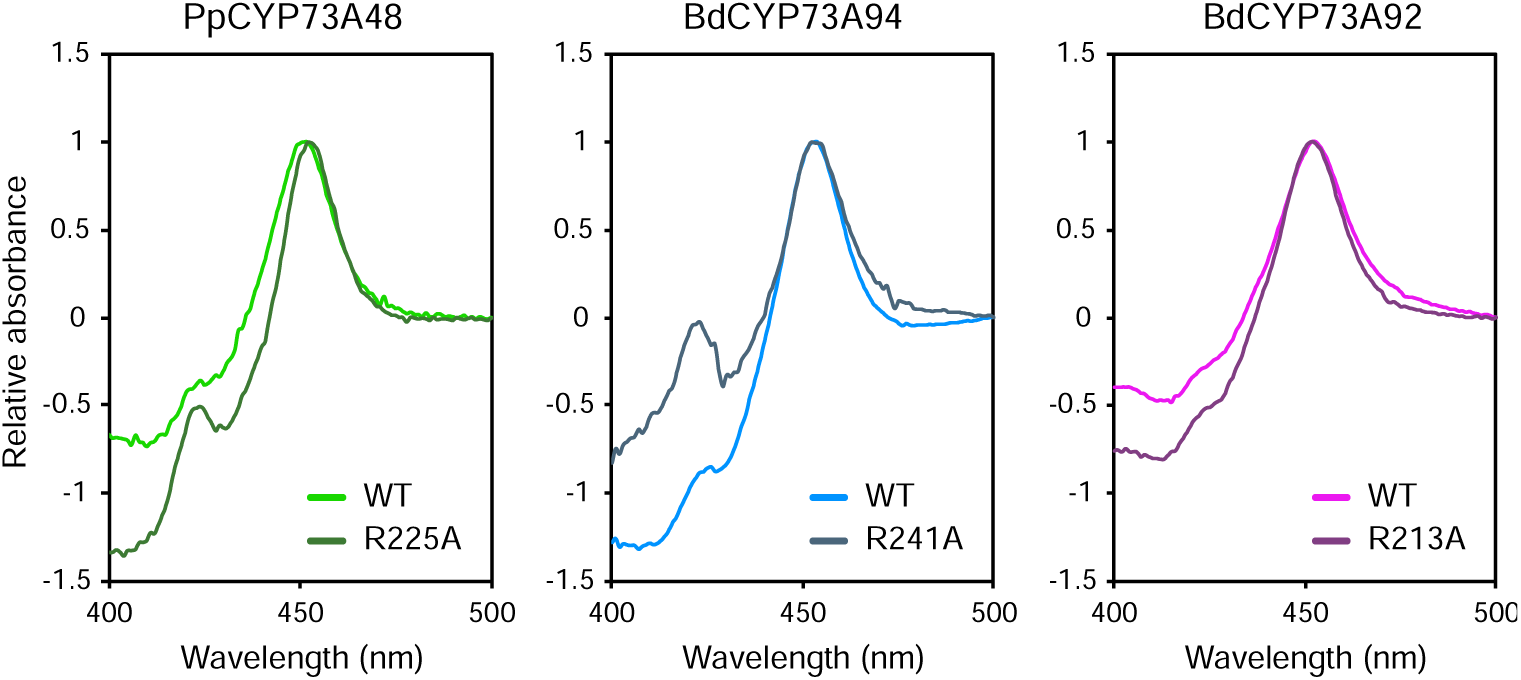
Cytochromes P450 CO difference spectra of wild-type and R>A mutated CYP73 proteins. CO difference spectra were determined from 20-fold dilution of microsomal preparations. Each absorbance spectrum was normalized according to its maximum.

**Figure S3.**
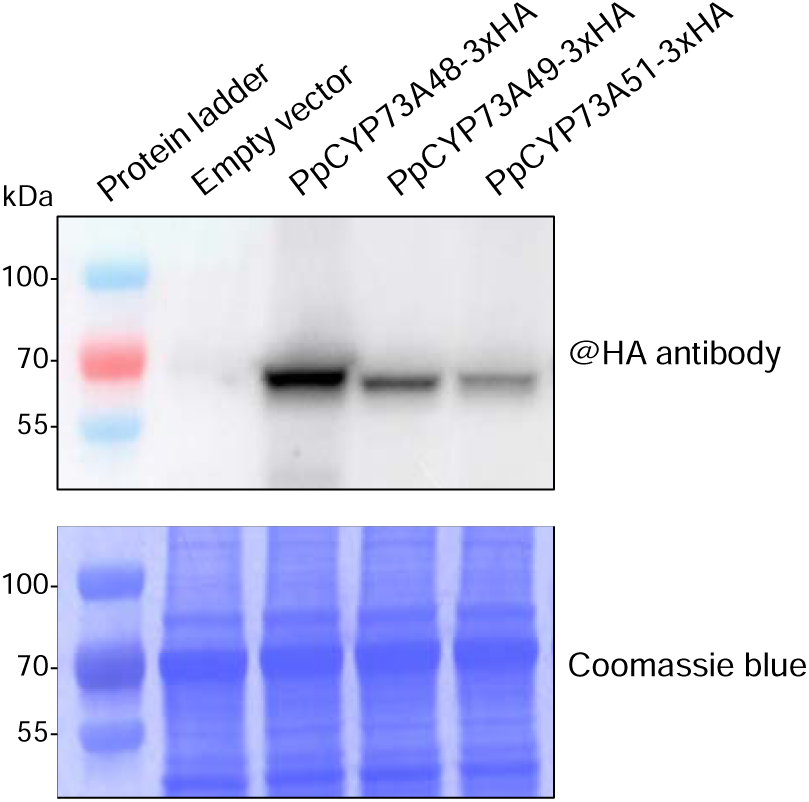
Western blot analysis of recombinant PpPpCYP73A-3xHA proteins. Microsomes were 10-fold diluted in denaturation buffer (200 mM Tris pH 6.8, 8% SDS, 40% glycerol, 0.1% bromophenol blue and 100 mM DTT) prior to denaturation at 65°C for 5 min. Ten micrograms total protein were loaded on each lane.

**Figure S4.**
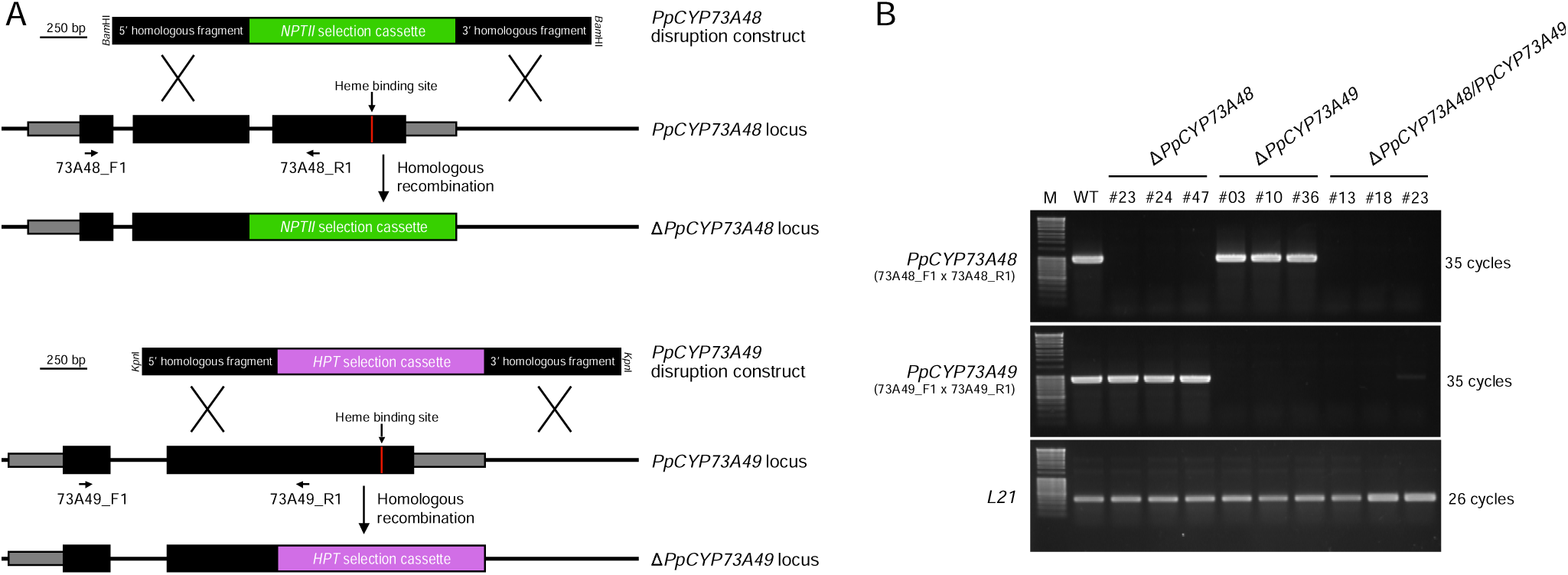
Molecular characterization of *Physcomitrium patens* Δ*CYP73A* mutants. (A) Homologous recombination-mediated strategy for *PpCYP73A48* and *PpCYP73A49* gene disruption. Genomic fragments encompassing the critical heme-binding site were excised with simultaneous insertion of the *NPTII* (*PpCYP73A48*) or *HPT* (*PpCYP73A49*) selection cassette conferring resistance to geneticin and hygromycin B, respectively. (B) RT-PCR analysis of Δ*PpCYP73A48,* Δ*PpCYP73A49* and Δ*PpCYP73A48/PpCYP73A49* mutant lines showing the absence of corresponding transcripts. The primer hybridization sites for RT-PCR are indicated in panel A. RT-PCR analysis of *L21* transcripts was used as amplification control. M, MassRuler™ DNA Ladder (ThermoFisher Scientific).

**Figure S5.**
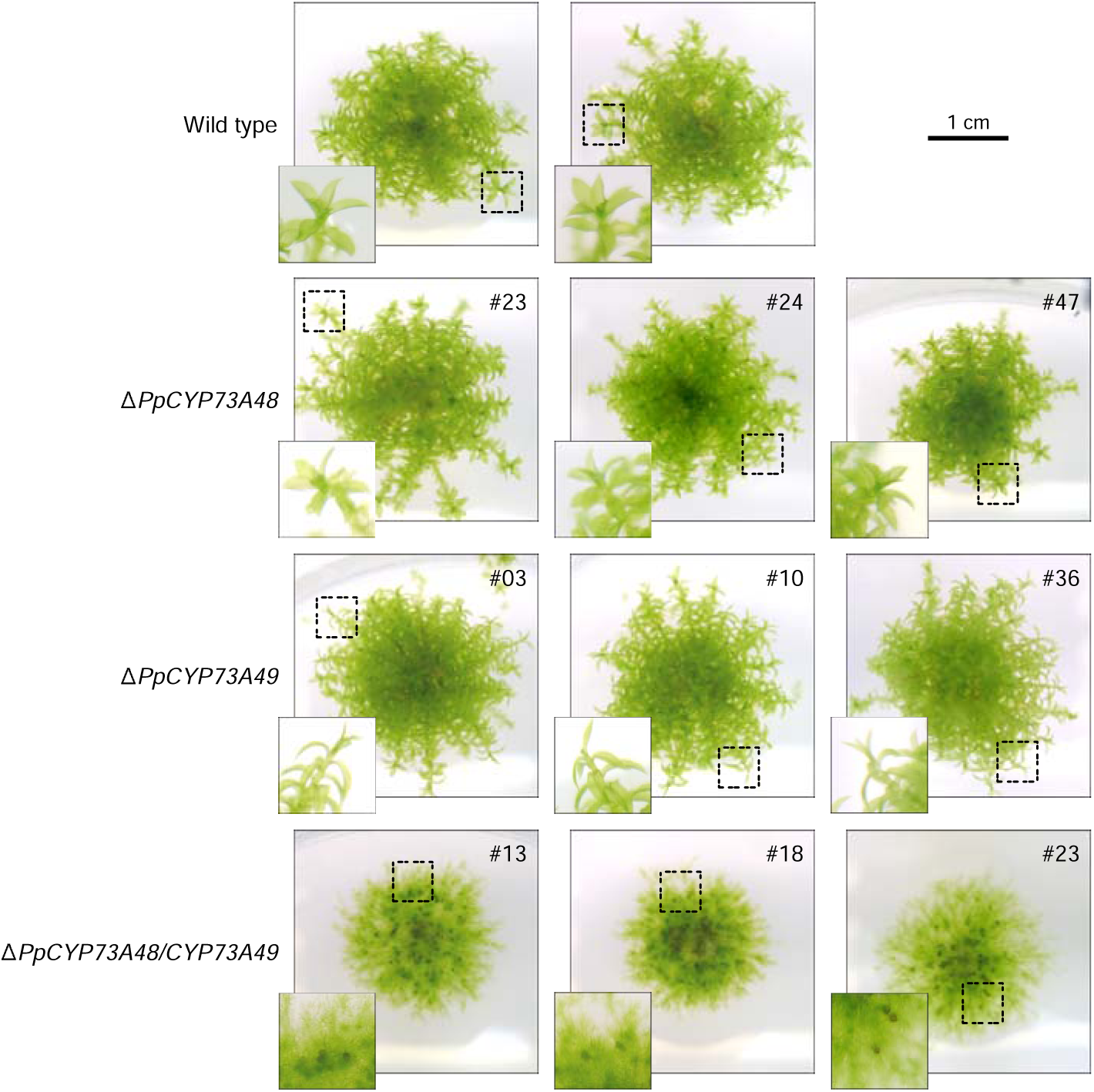
Macroscopic phenotypes of *Physcomitrium patens* Δ*CYP73A* mutants. Pictures of two-month-old *P. patens* colonies highlighting the consistent phenotypes observed across three independent lines for each genotype. For each colony, a 2.4x magnification of boxed area is shown.

**Figure S6.**
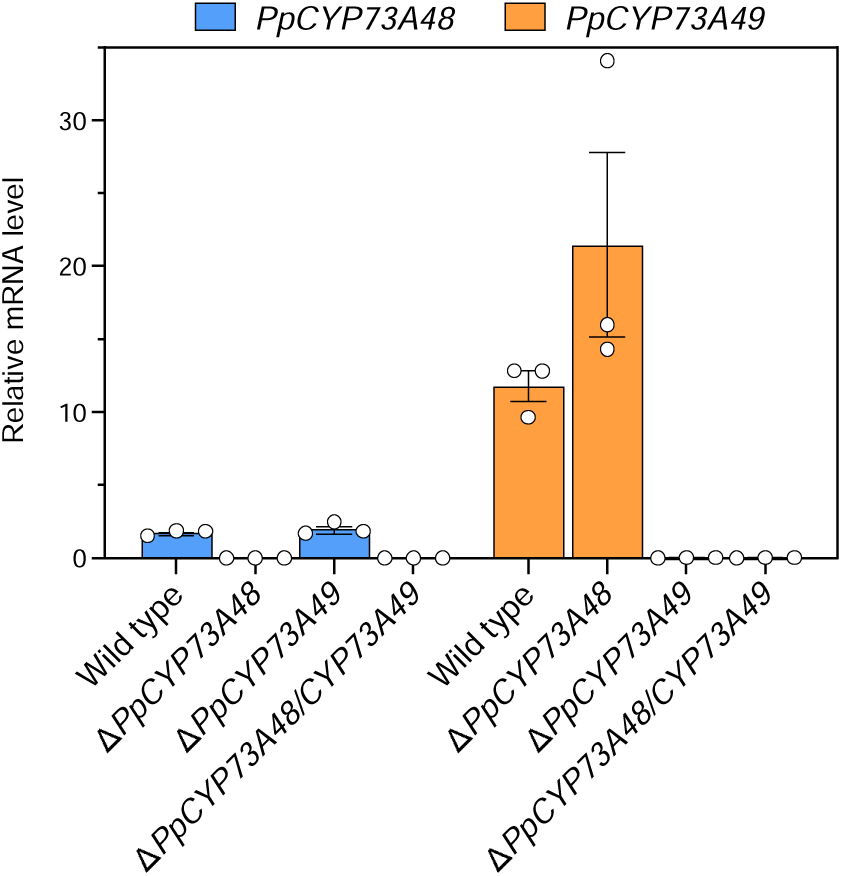
Expression analysis of *PpCYP73A48* and *PpCYP73A49* genes in mutant backgrounds. Relative mRNA levels were determined by RT-qPCR in wild type, Δ*PpCYP73A48,* Δ*PpCYP73A49*and Δ*PpCYP73A48/CYP73A49* mutant lines. Results are the mean ± SEM of three independent WT biological replicates and three independent mutant lines.

**Figure S7.**
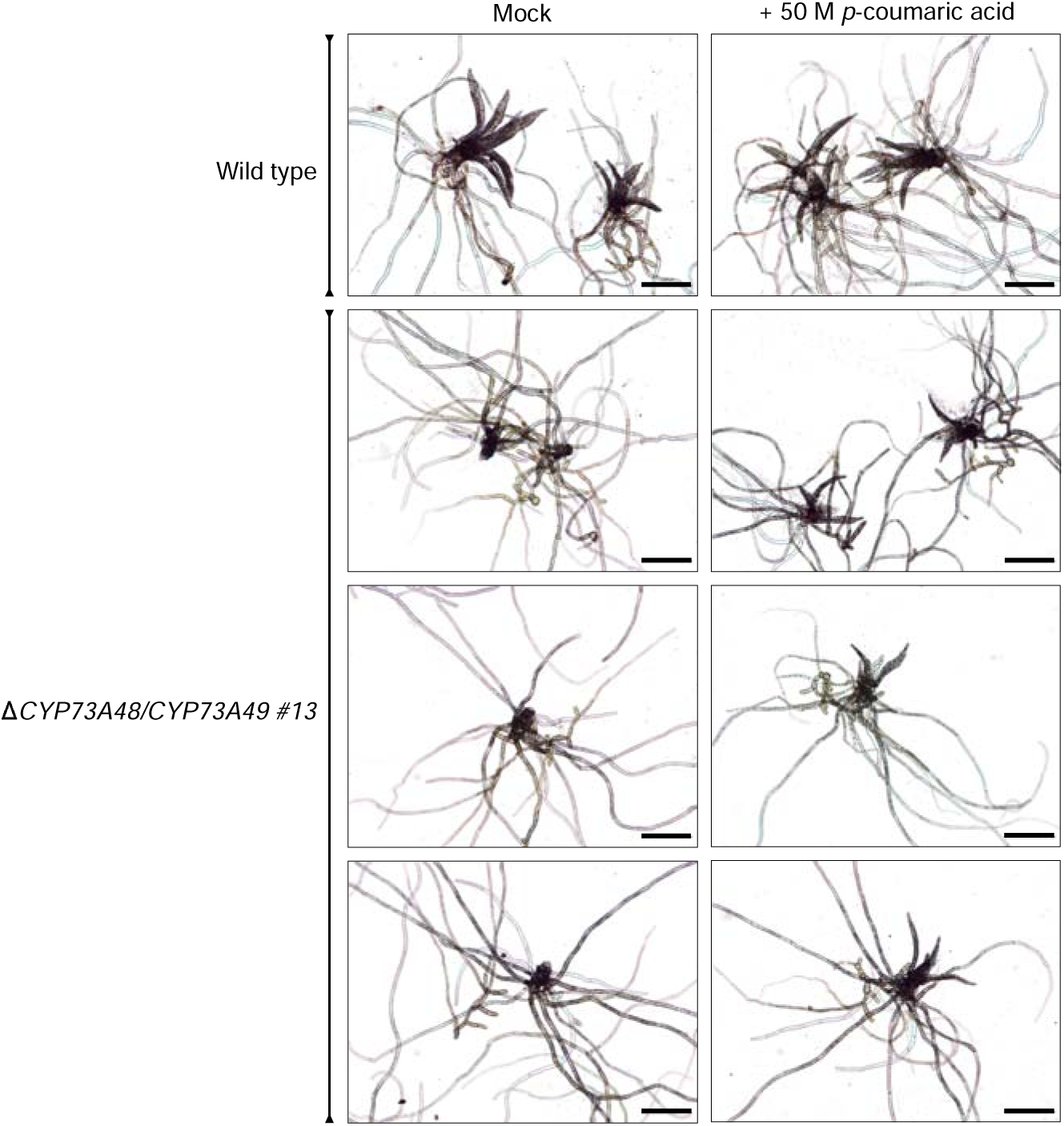
Chemical complementation of Δ*PpCYP73A48/CYP73A49* gametophore stunted growth with *p*-coumaric acid. Protoplasts from wild type and Δ*PpCYP73A48/CYP73A49* #13 mutant line were embedded in low-melting point agarose and regenerated in the presence of 50 µM *p*-coumaric acid or a corresponding mock medium (0.1% ethanol). Pictures were taken five weeks after start of regeneration and illustrate the positive effect of *p*-coumaric acid in rescuing the stunted growth of Δ*PpCYP73A48/CYP73A49* #13 gametophores. No significant effects of *p*-coumaric acid were observed in the wild type. Scale bars, 0.25 mm.

**Figure S8.**
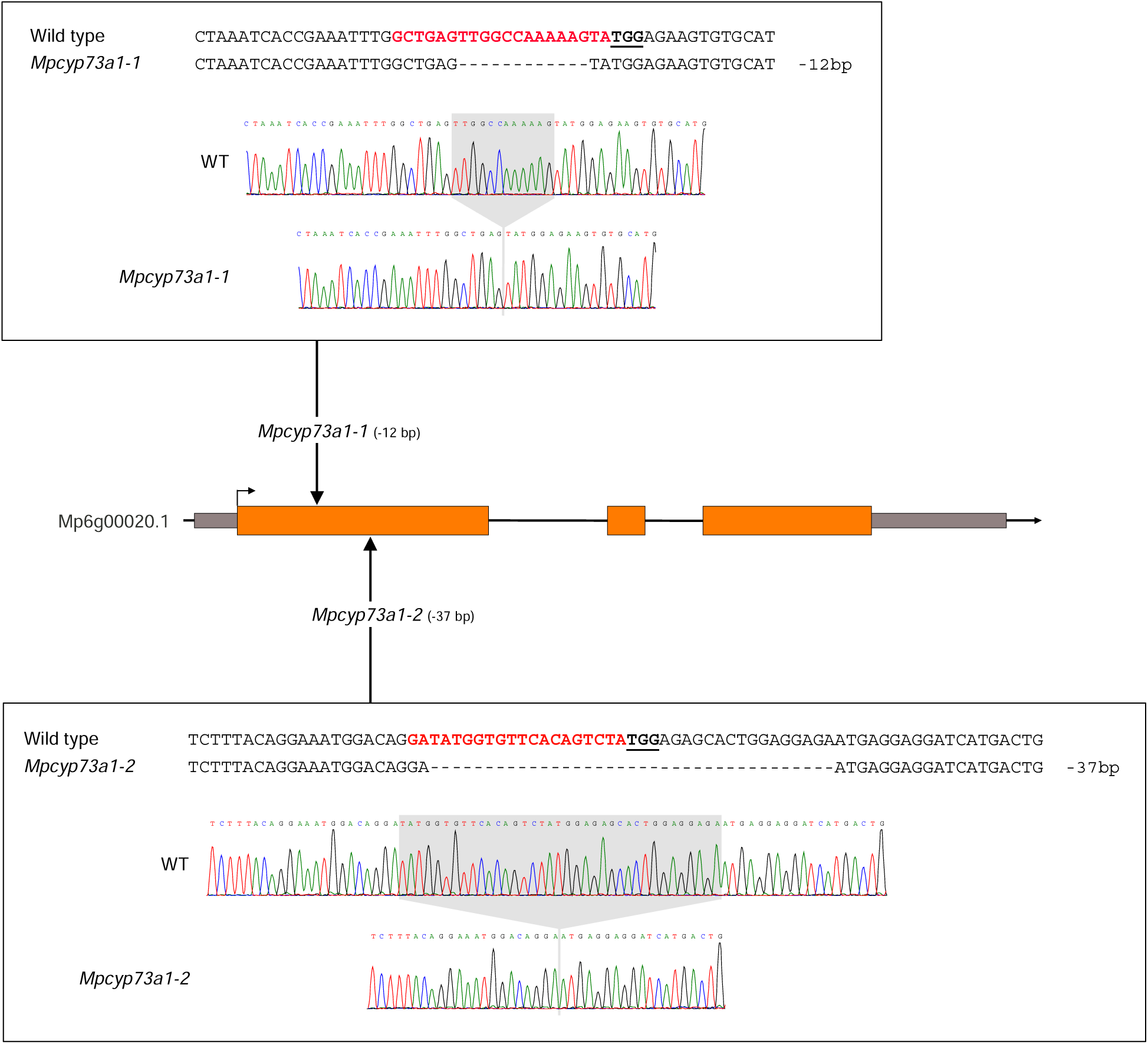
Molecular characterization of *Mpcyp73a1* CRISPR mutants. Two independent *Mpcyp73a1* CRISPR mutant lines were isolated and characterized by Sanger sequencing. The protospacer sequences, their corresponding locations, and the sequencing results are shown for both mutant lines. *Mpcyp73a1-1* exhibits a deletion of 12 nucleotides, while *Mpcyp73a1-2* displays a deletion of 37 nucleotides.

**Figure S9.**
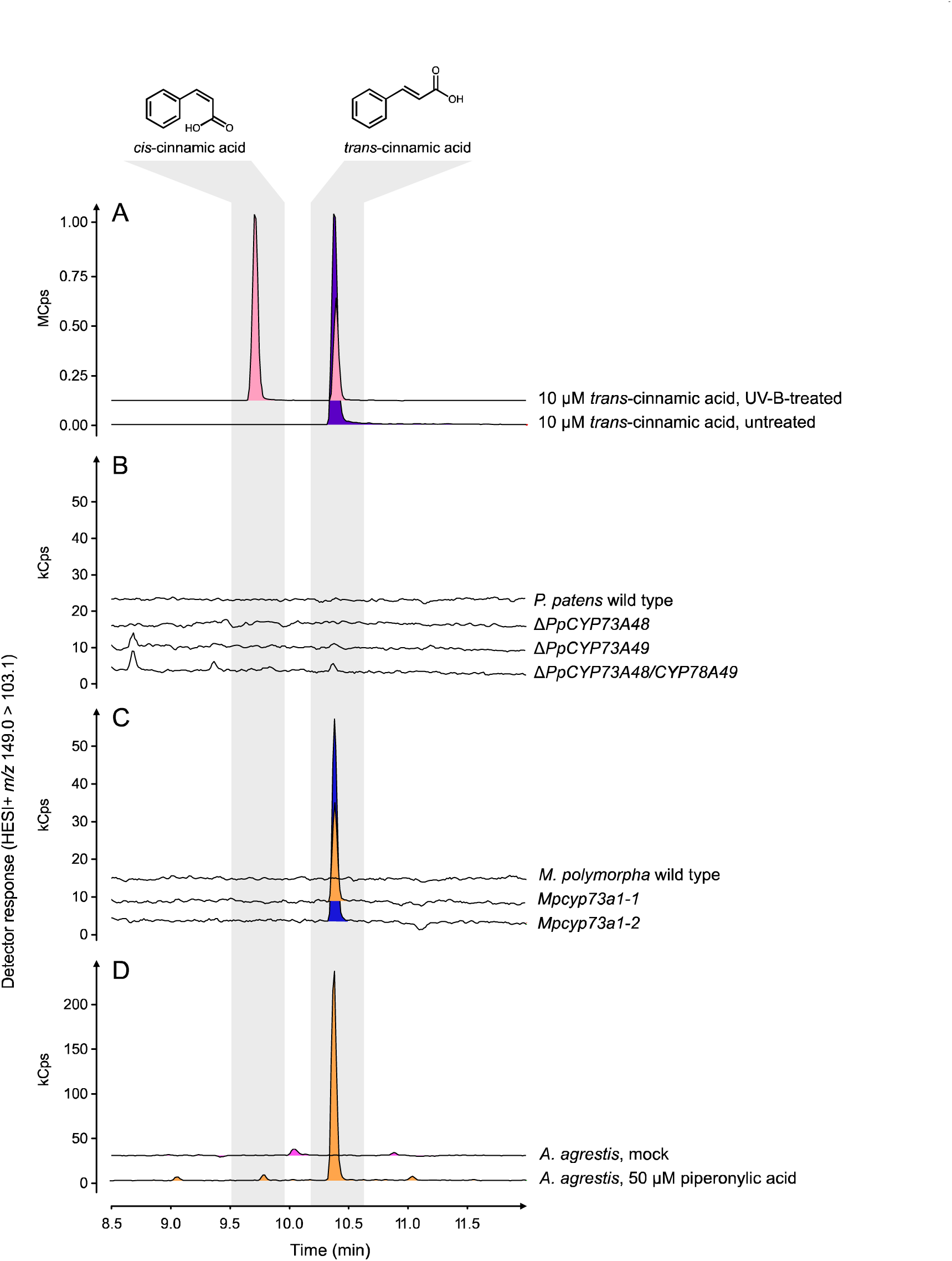
Search for *cis*-cinnamic acid in C4H-impaired plants. (A) A 10 µM *trans*-cinnamic acid solution was treated for 15 min with UV-B light provided by four UVB Broadband TL 40W/12 RS SLV tubes (Phillips), resulting in the appearance of the stereoisomer *cis*-cinnamic acid. Both isomers were then searched by UHPLC-MS/MS in crude extracts of (B) *P. patens* wild type and Δ*CYP73A* mutants (see Fig. 4), (C) *M. polymorpha* wild type and *Mpcyp73a1* CRISPR mutants (see Fig. 5), and in (D) *A. agrestis* treated with 50 µM piperonylic acid or corresponding mock treatment (see Fig. 6). Shown are representative UHPLC-MS/MS chromatograms.

**Table S2.**
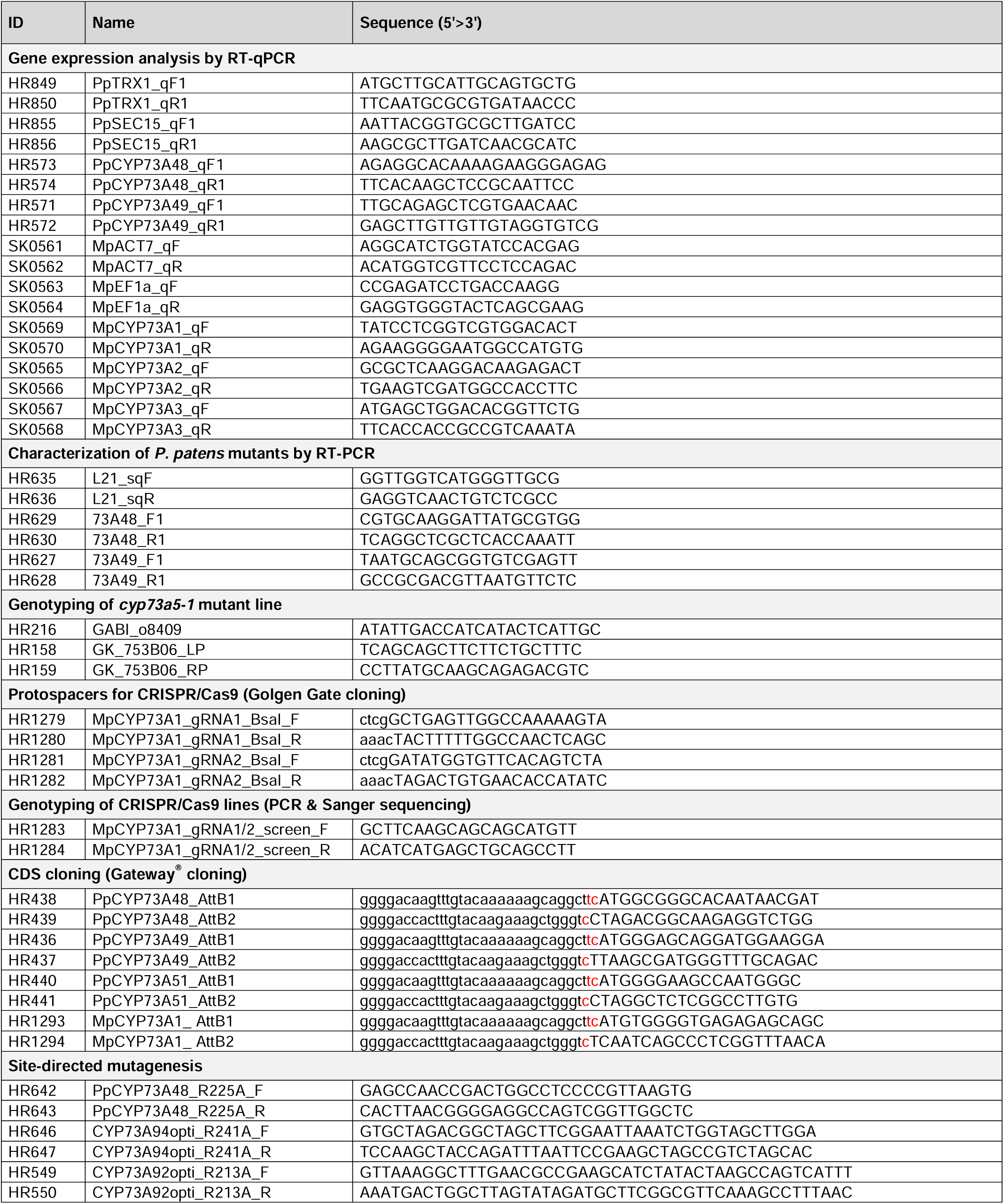
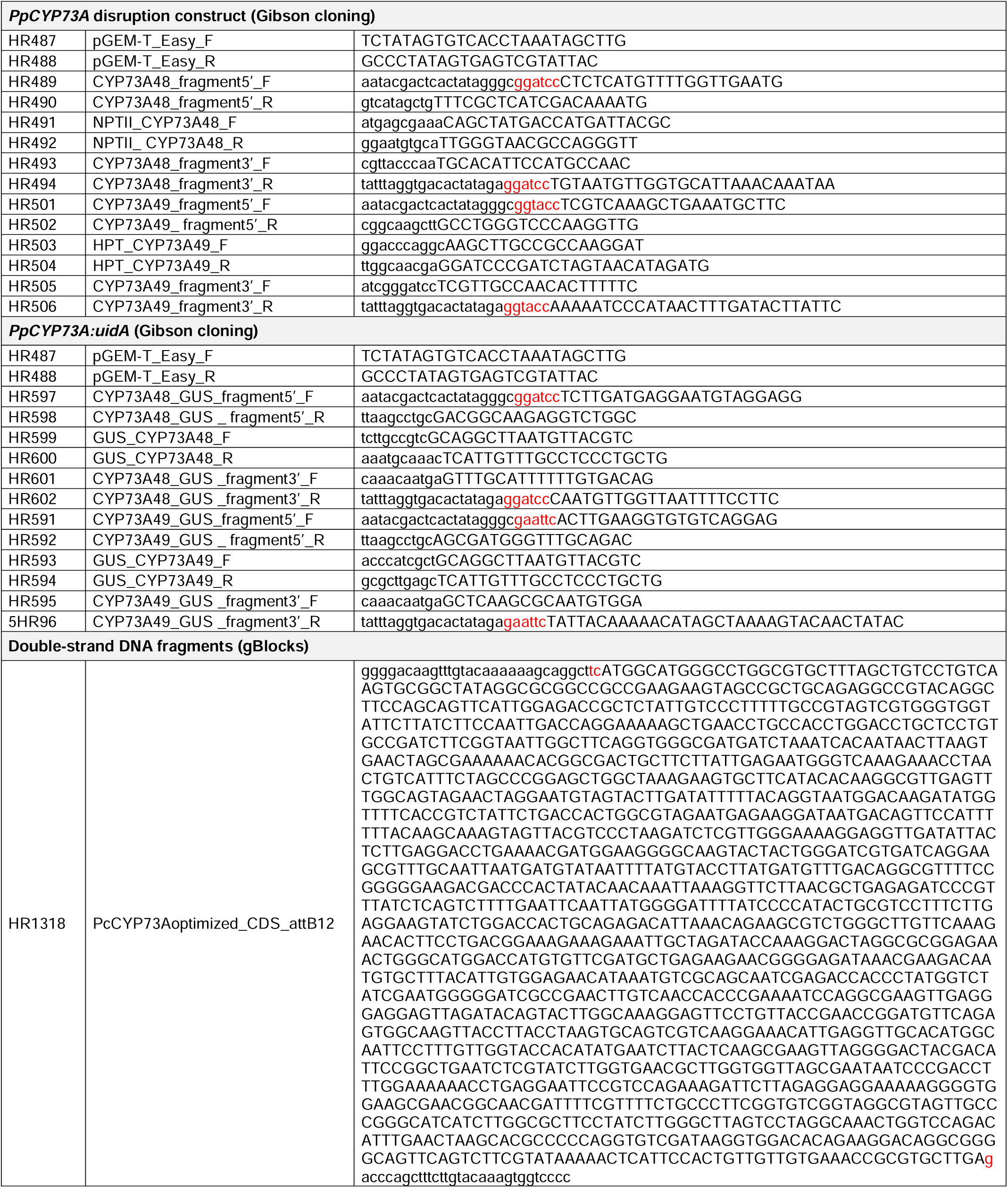
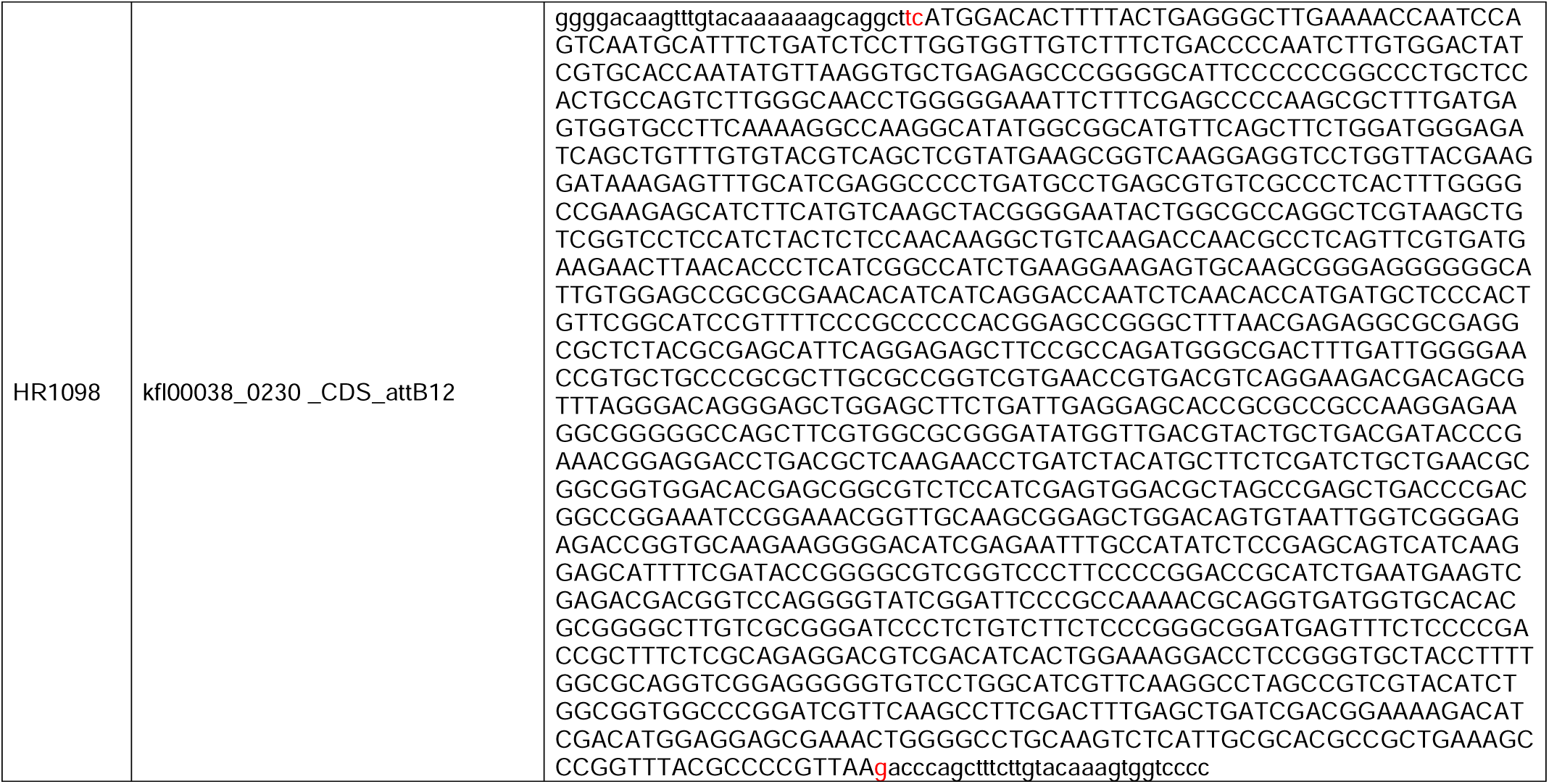
List of primers and synthesized sequences used in the study.

**Table S3.** List of multiple reaction monitoring (MRM) methods used for targeted analysis by UHPLC-MS/MS.

| Molecule | Ionization mode | Precursor ion | Collision energy | Product ion |
| --- | --- | --- | --- | --- |
| cinnamic acid | positive | 149.0 | 21.0 | 103.1 |
| *p*-coumaric acid | negative | 163.0 | 9.0 | 119.1 |
| caffeic acid | negative | 179.0 | 18.0 | 135.1 |
| ferulic acid | positive | 195.0 | 10.0 | 177.0 |
| *p*-coumaroyl-threonate | positive | 283.0 | 10.0 | 147.0 |
| *p*-coumaroyl-shikimate | positive | 320.9 | 8.0 | 147.0 |
| caffeoyl-threonate | positive | 299.0 | 38.0 | 89.0 |
| caffeoyl-shikimate | positive | 337.4 | 13.0 | 163.0 |
| rosmarinic acid | negative | 359.1 | 8.0 | 161.0 |

**Table S4.**
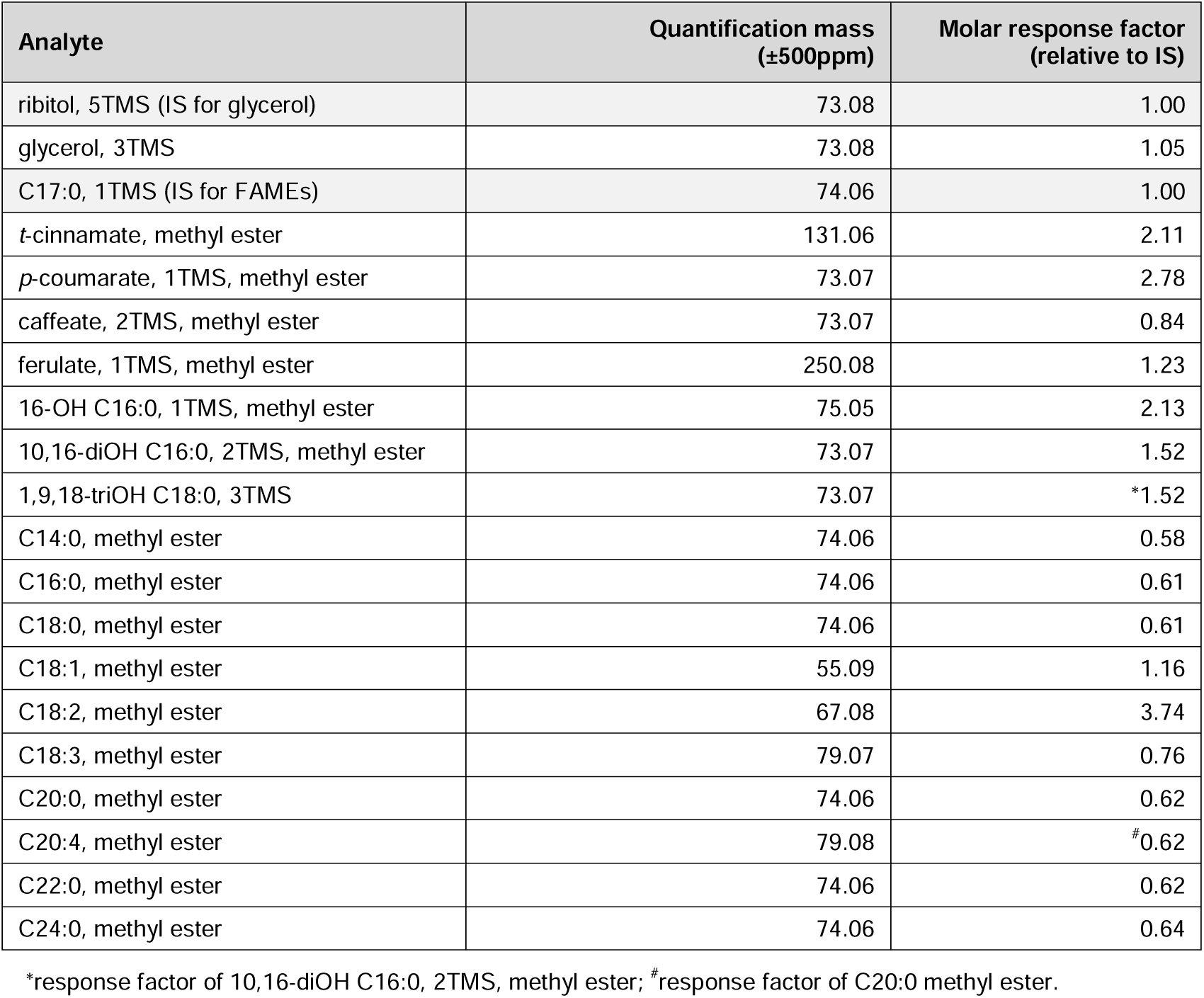
List of quantification masses and molar response factors used for quantitative analysis of cuticular monomers by GC-TOFMS. IS, internal standard; TMS, Trimethylsilyl group.

## Notes

### Competing Interest Statement

The authors have declared no competing interest.

## REFERENCES

1. Y. M. Bar-On, R. Phillips, R. Milo, The biomass distribution on Earth. Proc. Natl. Acad. Sci. USA 115, 6506– 6511 (2018).

2. S. Cheng, et al., Genomes of Subaerial Zygnematophyceae Provide Insights into Land Plant Evolution. Cell 179, 1057–1067.e14 (2019).

3. J. L. Morris, et al., The timescale of early land plant evolution. Proc. Natl. Acad. Sci. USA 115, 201719588 (2018).

4. T. Vogt, Phenylpropanoid biosynthesis. Molecular Plant 3, 2–20 (2010).

5. J. K. Weng, C. Chapple, The origin and evolution of lignin biosynthesis. New Phytologist 187, 273–285 (2010).

6. W. Boerjan, J. Ralph, M. Baucher, Lignin biosynthesis. Annu Rev Plant Biol 54, 519–546 (2003).

7. M. Mizutani, D. Ohta, R. Sato, Isolation of a cDNA and a Genomic Clone Encoding Cinnamate 4-Hydroxylase from Arabidopsis and Its Expression Manner in Planta. Plant Physiology 113, 755–763 (1997).

8. J.-E. Bassard, et al., Protein-protein and protein-membrane associations in the lignin pathway. Plant Cell 24, 4465–4482 (2012).

9. J. Barros, R. A. Dixon, Plant Phenylalanine/Tyrosine Ammonia-lyases. Trends in Plant Science 25, 66–79 (2019).

10. H. Renault, et al., Gene duplication leads to altered membrane topology of a cytochrome P450 enzyme in seed plants. Mol Biol Evol 34, 2041–2056 (2017).

11. B. Hamberger, S. Bak, Plant P450s as versatile drivers for evolution of species-specific chemical diversity. Philos Trans R Soc Lond B Biol Sci 368, 20120426 (2013).

12. Z. Liu, et al., Evolutionary interplay between sister cytochrome P450 genes shapes plasticity in plant metabolism. Nature Communications 7, 13026 (2016).

13. C. C. Hansen, D. R. Nelson, B. L. Møller, D. Werck-Reichhart, Plant cytochrome P450 plasticity and evolution. Molecular Plant 14, 1244–1265 (2021).

14. A. L. Schilmiller, et al., Mutations in the cinnamate 4-hydroxylase gene impact metabolism, growth and development in Arabidopsis. Plant Journal 60, 771–782 (2009).

15. R. Hu, et al., Adaptive evolution of the enigmatic Takakia now facing climate change in Tibet. Cell 186, 3558– 3576 (2023).

16. B. Zhang, et al., Structure and Function of the Cytochrome P450 Monooxygenase Cinnamate 4-hydroxylase from Sorghum bicolor. Plant Physiology 183, 957–973 (2020).

17. H. Renault, et al., A phenol-enriched cuticle is ancestral to lignin evolution in land plants. Nature Communications 8, 14713 (2017).

18. L. Kriegshauser, et al., Function of the HYDROXYCINNAMOYL-CoA:SHIKIMATE HYDROXYCINNAMOYL TRANSFERASE is evolutionarily conserved in embryophytes. Plant Cell 33, 1472–1491 (2021).

19. H. Berland, et al., Auronidins are a previously unreported class of flavonoid pigments that challenges when anthocyanin biosynthesis evolved in plants. Proc. Natl. Acad. Sci. USA 116, 20232–20239 (2019).

20. M. Schalk, et al., Piperonylic acid, a selective, mechanism-based inactivator of the trans-cinnamate 4-hydroxylase: A new tool to control the flux of metabolites in the phenylpropanoid pathway. Plant Physiology 118, 209–218 (1998).

21. S. Naseer, et al., Casparian strip diffusion barrier in Arabidopsis is made of a lignin polymer without suberin. Proc. Natl. Acad. Sci. USA 109, 10101–10106 (2012).

22. J. Wohl, M. Petersen, Functional expression and characterization of cinnamic acid 4-hydroxylase from the hornwort Anthoceros agrestis in Physcomitrella patens. Plant Cell Rep 39, 597–607 (2020).

23. M. Petersen, et al., Evolution of rosmarinic acid biosynthesis. Phytochemistry 70, 1663–1679 (2009).

24. I. E. Houari, et al., Seedling developmental defects upon blocking CINNAMATE-4-HYDROXYLASE are caused by perturbations in auxin transport. New Phytologist 230, 2275–2291 (2021).

25. Sarkanen, K. V., Ludwig, C. H., Lignins.J: occurrence, formation, structure and reactions, (John Wiley & Sons, Inc., New York, 1971).

26. E. Sjöström, Wood Chemistry Fundamentals and Applications, (Academic Press Inc, 1993).

27. J. de Vries, S. de Vries, C. H. Slamovits, L. E. Rose, J. M. Archibald, How embryophytic is the biosynthesis of phenylpropanoids and their derivatives in streptophyte algae? Plant and Cell Physiology 58, 934–945 (2017).

28. S. de Vries, et al., The evolution of the phenylpropanoid pathway entailed pronounced radiations and divergences of enzyme families. Plant Journal 107, 975–1002 (2021).

29. H. Renault, D. Werck-Reichhart, J.-K. Weng, Harnessing lignin evolution for biotechnological applications. Current Opinion in Biotechnology 56, 105–111 (2019).

30. M. Mizutani, D. Ohta, Diversification of P450 genes during land plant evolution. Annu Rev Plant Biol 61, 291– 315 (2010).

31. J. Barros, et al., Role of bifunctional ammonia-lyase in grass cell wall biosynthesis. Nature Plants 2, 16050 (2016).

32. S. Nedelkina, et al., Novel characteristics and regulation of a divergent cinnamate 4-hydroxylase (CYP73A15) from French bean: engineering expression in yeast. Plant Mol Biol 39, 1079–1090 (1999).

33. M. A. Pierrel, et al., Catalytic properties of the plant cytochrome P450 CYP73 expressed in yeast. Substrate specificity of a cinnamate hydroxylase. European Journal of Biochemistry 224, 835–844 (1994).

34. Y.-F. Wu, et al., Isolation and functional characterization of hydroxycinnamoyltransferases from the liverworts Plagiochasma appendiculatum and Marchantia paleacea. Plant Physiology and Biochemistry 129, 400–410 (2018).

35. M. Schalk, et al., Design of fluorescent substrates and potent inhibitors of CYP73As, P450s that catalyze 4-hydroxylation of cinnamic acid in higher plants. Biochemistry 36, 15253–15261 (1997).

36. H. Chen, H. Jiang, J. A. Morgan, Non-natural cinnamic acid derivatives as substrates of cinnamate 4-hydroxylase. Phytochemistry 68, 306–311 (2007).

37. L. Achnine, E. B. Blancaflor, S. Rasmussen, R. A. Dixon, Colocalization of L-phenylalanine ammonia-lyase and cinnamate 4-hydroxylase for metabolic channeling in phenylpropanoid biosynthesis. Plant Cell 16, 3098– 3109 (2004).

38. R. Franke, et al., Changes in secondary metabolism and deposition of an unusual lignin in the ref8 mutant of Arabidopsis. Plant journal 30, 47–59 (2002).

39. L. Hoffmann, et al., Silencing of hydroxycinnamoy-coenzyme A shikimate/quinate hydroxycinnamoyltransferase affects phenylpropanoid biosynthesis. Plant Cell 16, 1446–1465 (2004).

40. J. Huang, et al., Functional analysis of the Arabidopsis PAL gene family in plant growth, development, and response to environmental stress. Plant Physiology 153, 1526–1538 (2010).

41. Y. Li, J. I. Kim, L. Pysh, C. Chapple, Four isoforms of Arabidopsis 4-Coumarate:CoA ligase have overlapping yet distinct roles in phenylpropanoid metabolism. Plant Physiology 169, 2409–2421 (2015).

42. F. Muro-Villanueva, X. Mao, C. Chapple, Linking phenylpropanoid metabolism, lignin deposition, and plant growth inhibition. Current Opinion in Biotechnology 56, 202–208 (2019).

43. J. H. Leebens-Mack, et al., One thousand plant transcriptomes and the phylogenomics of green plants. Nature 574, 679–685 (2019).

44. R. C. Edgar, MUSCLE: multiple sequence alignment with high accuracy and high throughput. Nucleic Acids Res 32, 1792–1797 (2004).

45. J. Castresana, Selection of conserved blocks from multiple alignments for their use in phylogenetic analysis. Mol Biol Evol 17, 540–552 (2000).

46. M. Gouy, S. Guindon, O. Gascuel, SeaView version 4: A multiplatform graphical user interface for sequence alignment and phylogenetic tree building. Mol Biol Evol 27, 221–224 (2010).

47. B. Q. Minh, et al., IQ-TREE 2: New Models and Efficient Methods for Phylogenetic Inference in the Genomic Era. Mol Biol Evol 37, 1530–1534 (2020).

48. A. Waterhouse, et al., SWISS-MODEL: homology modelling of protein structures and complexes. Nucleic Acids Research 46, W296–W303 (2018).

49. O. Trott, A. J. Olson, AutoDock Vina: Improving the speed and accuracy of docking with a new scoring function, efficient optimization, and multithreading. J Comput Chem 31, 455–461 (2010).

50. E. F. Pettersen, et al., UCSF ChimeraX: Structure visualization for researchers, educators, and developers. Protein Science 30, 70–82 (2021).

51. S. Alberti, A. D. Gitler, S. Lindquist, A suite of Gateway® cloning vectors for high-throughput genetic analysis in Saccharomyces cerevisiae. Yeast 24, 913–919 (2007).

52. P. Urban, C. Mignotte, M. Kazmaier, F. Delorme, D. Pompon, Cloning, yeast expression, and characterization of the coupling of two distantly related Arabidopsis thaliana NADPH-cytochrome P450 reductases with P450 CYP73A5. J Biol Chem 272, 19176–19186 (1997).

53. F. P. Guengerich, M. V. Martin, C. D. Sohl, Q. Cheng, Measurement of cytochrome P450 and NADPH– cytochrome P450 reductase. Nature Protocols 4, 1245–1251 (2009).

54. S. A. Rensing, et al., The Physcomitrella genome reveals evolutionary insights into the conquest of land by plants. Science 319, 64–69 (2008).

55. F. W. Li, et al., Anthoceros genomes illuminate the origin of land plants and the unique biology of hornworts. Nature Plants 6, 259–272 (2020).

56. J. L. Bowman, et al., Insights into land plant evolution garnered from the Marchantia polymorpha genome. Cell 171, 287–304 (2017).

57. T. Girke, H. Schmidt, U. Zahringer, R. Reski, E. Heinz, Identification of a novel delta 6-acyl-group desaturase by targeted gene disruption in Physcomitrella patens. Plant Journal 15, 39–48 (1998).

58. J. K. Weng, T. Akiyama, J. Ralph, C. Chapple, Independent recruitment of an O-methyltransferase for syringyl lignin biosynthesis in Selaginella moellendorffii. Plant Cell 23, 2708–2724 (2011).

59. S. J. Clough, A. F. Bent, Floral dip: A simplified method for Agrobacterium-mediated transformation of Arabidopsis thaliana. Plant Journal 16, 735–743 (1998).

60. G. Wiedemann, et al., RecQ Helicases Function in Development, DNA Repair, and Gene Targeting in Physcomitrella patens. Plant Cell 30, 717–736 (2018).

61. G. Philippe, et al., Ester cross-link profiling of the cutin polymer of wild-type and cutin synthase tomato mutants highlights different mechanisms of polymerization. Plant Physiology 170, 807–20 (2016).

62. S. S. Sugano, et al., Efficient CRISPR/Cas9-based genome editing and its application to conditional genetic analysis in Marchantia polymorpha. PloS One 13, e0205117 (2018).

63. J.-P. Concordet, M. Haeussler, CRISPOR: intuitive guide selection for CRISPR/Cas9 genome editing experiments and screens. Nucleic Acids Research 46, W242–W245 (2018).

64. A. Kubota, K. Ishizaki, M. Hosaka, T. Kohchi, Efficient Agrobacterium-mediated transformation of the liverwort Marchantia polymorpha using regenerating thalli. Bioscience, Biotechnology, and Biochemistry 77, 167–172 (2013).

65. S. Proost, M. Mutwil, CoNekT: an open-source framework for comparative genomic and transcriptomic network analyses. Nucleic Acids Research 46, W133–W140 (2018).

